# Determinants of metal import and specificity in a bacterial transporter

**DOI:** 10.64898/2026.03.30.714904

**Authors:** Samuel P. Berry, Camille B. Freedman, Debora S. Marks, Rachelle Gaudet

## Abstract

Membrane transporters have evolved finely tuned substrate specificity within a limited repertoire of structural superfamilies, but the biophysical origins of transporter specificity remain unclear. We have systematically investigated the determinants of import and specificity in the model *Deinococcus radiodurans* (Dra)Nramp metal importer by testing targeted structure- and evolution-guided libraries of sequence variants in new high-throughput assays for import of the native substrate Mn^2+^ and the chemically similar but excluded metal Mg^2+^. The effects of most combinatorial mutations on Mn^2+^ import fit a simple global epistasis model, but many additional unexplained long-range epistatic interactions cluster at a set of structural hotspots at the inner and outer vestibules of the transporter. By contrast, mutations enabling Mg^2+^ import have non-additive effects and generally fall into two categories: a few core specificity positions in the first two shells around the metal are sufficient to allow DraNramp to import Mg^2+^, and additional modulator mutations can finetune specificity when combined with core-position mutations. The modulator mutations overlap considerably with key epistatic mutations, and we use this insight to propose a new biochemical model for how mutations that alter conformational balance in transporters can lead both to long-range epistasis and specificity modulation.

## Introduction

Membrane transport proteins control what molecules enter and exit the cell based on their ability to accurately discriminate between chemical substrates. Most transporters come from a small number of structural superfamilies in which homologs with similar architecture enforce vastly different substrate specificities. The ability to do so is encoded in each transporter’s amino acid sequence, yet our ability to predict transporter specificity from biochemical first principles remains out of reach. Detailed structural studies of individual transporters can reveal how those transporters bind a given substrate, but they do not uncover the rules of substrate specificity within a transporter family, including how a given transporter excludes other substrates.

The Nramp (natural resistance associated macrophage protein) family of metal ion importers poses a chemically simple but puzzling example of such a specificity mandate: Nramps must be able to import scarce divalent transition metals like Mn^2+^ and Fe^2+^ while excluding far more abundant alkaline earth metals such as Mg^2+^ and Ca^2+^ ^1,2^. Import of scarce transition metals is essential for various metabolic processes in all living cells, and imbalances in cellular concentrations of transition metals are linked to range of disorders including anemia and neurodegenerative diseases^3^. Canonical Nramps, found from bacteria to humans, transport transition metals with little to no inhibition by alkaline earth metals, despite their much higher concentrations in typical biological solutions^2,4,5^. This strict selectivity is remarkable given the chemical similarities between transition and alkaline earth metals. For example, Mn^2+^ and Mg^2+^ share the same charge (+2), preferred coordination geometry (octahedral), and typical bond lengths (2.1-2.5 Å)^6,7^.

Nramps present a compelling model for investigating the molecular mechanisms of transporter selectivity both because of this simple yet puzzling specificity as well as their extensive prior biochemical and structural characterization. Structures of a model bacterial Nramp from *Deinococcus radiodurans* (DraNramp) in multiple conformations with and without Mn^2+^ reveal how it binds a nearly dehydrated divalent cation via distinct coordination geometries in each conformational state^8^. All these geometries deviate from the ideal Mn^2+^ coordination geometry seen in high-affinity metal-binding proteins^6,9^, which suggests that overly tight binding in a transporter would inhibit transport. The same set of amino acid sidechains—primarily an aspartate and asparagine on transmembrane helix 1 (TM1; D56 and N59 in DraNramp) and a methionine on TM6 (M230A in DraNramp)—form a metal-binding motif strictly conserved across the canonical Nramps^8^. Nramps co-transport protons through a pathway partly distinct from that of the metal; this proton cotransport is not strictly required for metal import at high membrane potential but likely imparts a kinetic bias to favor import and allow for metal accumulation^10^.

While DraNramp’s specificity mandate is shared by canonical Nramps, more distantly related bacterial homologs have been identified with altered substrate specificity. Notably, an Nramp-related transporter from *Eggerthella lenta* (EleNRMT) transports Mg^2+^ nearly as well as Mn^2+^, likely transporting a more hydrated ion in its enlarged binding site^11^. While substituting the full set of binding-site residues from EleNRMT into the canonical bacterial *Eremococcus coleocola* Nramp (EcoDMT) did not enable Mg^2+^ transport^11^, the M230A substitution in DraNramp allowed for some Mg^2+^ import while still permitting Mn^2+^ import^1^. These results demonstrate that minor sequence perturbations can alter specificity to allow for Mg^2+^ import but also suggest that residues elsewhere in the protein likely play a role.

The are several possible explanations for how the binding-site methionine excludes Mg^2+^. Removing the methionine sidechain enlarges the binding site, allowing transport of a more hydrated ion, which is key for Mg^2+^ selectivity in channels^12^. For example, the Mg^2+^ channel CorA transports a partially hydrated Mg^2+^ ion while excluding transition metals by forming a selectivity filter too small for a fully hydrated Mn^2+^ or Fe^2+^ ion but too large for a dehydrated one^13,14^. By contrast, the occluded intermediate state of the DraNramp transport cycle requires stripping the Mn^2+^ ion of all but one of the six waters^8^. Additionally, Mg^2+^ is a “hard” Lewis acid, and thus it interacts less favorably with soft bases like the thioester of methionine compared to the “softer” acids Mn^2+^ and Fe^2+^ ^15^. However, this single example of a specificity-controlling position does not alone explain which of these factors are necessary or sufficient to alter specificity. Are there many routes to achieving altered metal specificity? Perhaps any change that enlarges the binding site to accommodate additional waters can increase Mg^2+^ transport? Do specificity-altering mutations always involve positions directly contacting the metal, or do allosteric mutations also play a role?

Previous studies on Nramps have tested individual mutations in low throughput^1,4,5,10,11,16,17^, limiting the ability to address these questions in a systematic manner. In recent years, however, approaches for testing thousands of mutations in parallel^18^ alongside novel machine learning methods based on natural sequence information enabled much more systematic profiling of the mutational landscape of proteins. Several studies profiling multiple substrates have highlighted how mutations distal to the binding site can affect substrate selectivity^19-21^, most relevantly for the human serotonin transporter^22^, a distant homolog of Nramps. Further, while most studies examine only single mutations, profiling energetics interactions between mutations—epistasis^23^—requires testing multiple mutations at once. However, because it is not possible to systematically profile all combinations of mutations, previous studies have used preexisting knowledge to narrow down the space of possibilities. Some studies condition based on structural intuition, systematically testing all combinations of mutations at a few defined positions such as the active site^24,25^. Others leverage natural sequence diversity either by sampling differences between known sequences^26,27^ or sampling directly from a computational model trained on natural sequences^28,29^. Some also use supervised models fit to preexisting mutational scanning data^30^.

In this work, we use principles from all three of the above sampling strategies to design a DraNramp library containing both a wide range of single variants and higher-order combinatorial variants. We develop assays amenable to screening this DraNramp library for metal import in high throughput to probe the fitness landscapes of DraNramp for both a native and non-native substrate and statistical models to analyze and interpret the effect of mutations. By accounting for the alternating-access requirement for substrate transport, we find that mutations with long-range epistasis and specificity modulation phenotypes likely alter the conformational balance of the transporter.

## Results

### Designing a mutational scanning library guided by structure and natural sequence diversity

To generate a mutational scanning library enriched for variants that alter Nramp metal specificity, we took two complementary approaches based on distinct hypotheses. First, single mutations such as M230A may be located at residues near the binding site, and some may have an observable effect only on the M230A background. We therefore designed a binding-site library of all single point mutations at DraNramp residues within a 12-Å shell of the orthosteric metal-binding site on both the wildtype (WT) and M230A background, totaling 2,900 variants (Fig. 1a; synthesis errors added 9,405 additional variants to the library).

**Figure 1.**
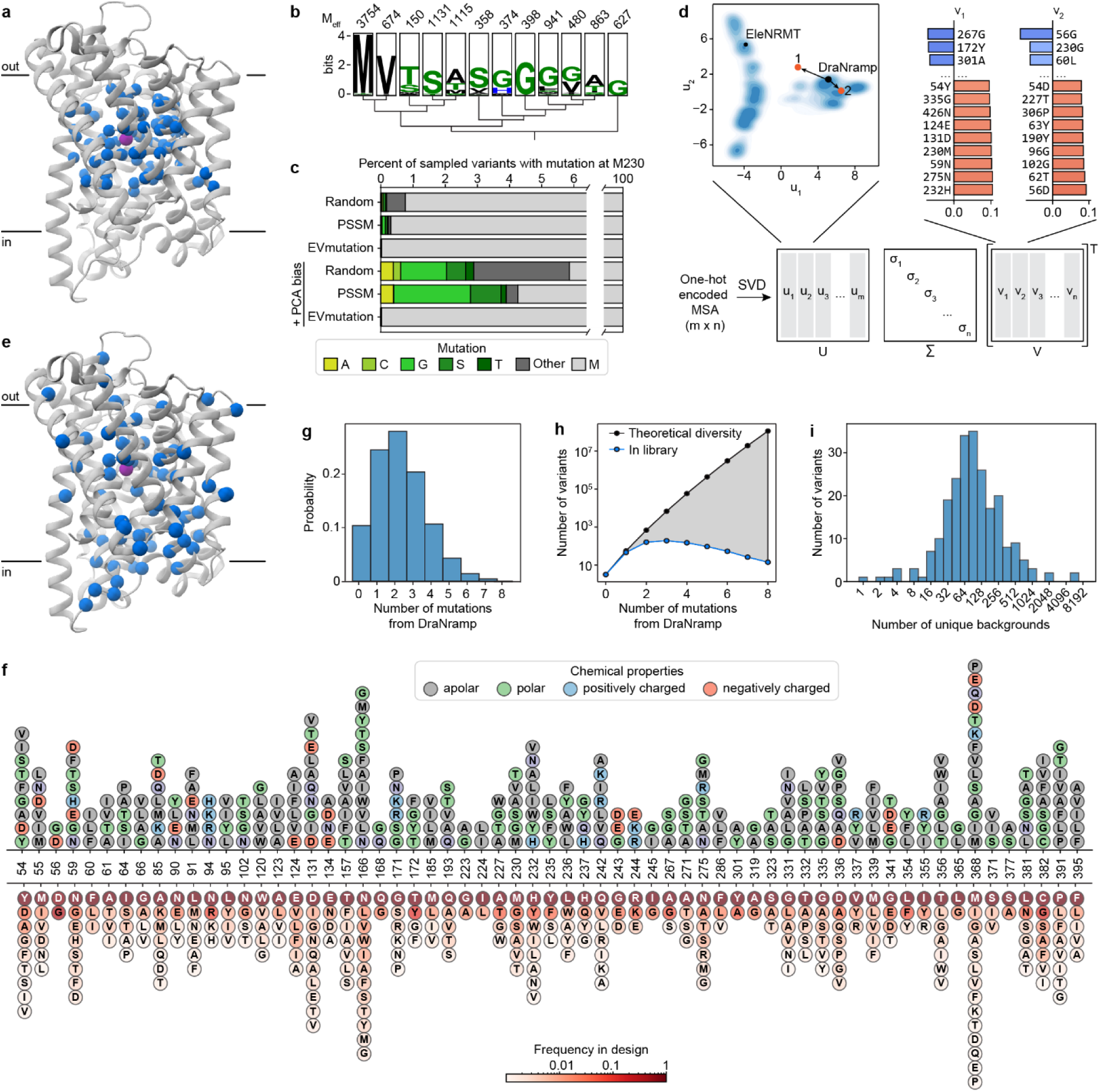
Design of a variant library leveraging structural and evolutionary information. (a) Blue spheres mark all positions within 12 Å of the metal-binding site in one of the three major conformations of DraNramp, which we mutated in the binding-site library; residues are highlighted on the Mn^2+^-bound occluded structure (PDB ID: 8E60), with Mn^2+^ shown in magenta. (b) Sequence logo showing the most abundant residue at the position aligned to M230 in DraNramp in canonical Nramps (left) and Nramp-related clades, in an MSA of 36,700 Nramp-related proteins^66^. The y-axis represents information in bits. The *M_eff_* values for each clade are indicated at the top. (c) The average frequency with which different sequence models trained on this diverse multiple sequence alignment sample known specificity-altering mutations at position 230 in DraNramp. In each case, models were restrained to sample variants with an average Hamming distance of approximately 3 from WT, and as such all models sample primarily methionine (the wildtype amino acid) at this position. (d) Schematic illustration of PCA on a one-hot encoded MSA using singular value decomposition (SVD). The upper left shows the resulting embedding vectors for the Nramp alignment. The positions of DraNramp and EleNRMT are marked. The upper right shows the values of the largest and smallest weights in the first two principal component vectors. These weights represent individual amino acids being at particular positions. (e) Blue spheres mark positions mutated in the final evolution-guided library design. (f) Mutations included in the design of the evolution-guided library, colored by chemical properties (top) or by frequency in the library (bottom). (g) Probability distribution of variants with different given numbers of mutations in the evolution-guided library design. (h) Comparison of the observed number of variants across mutational depths in the synthesized library (blue) against the theoretical diversity of the evolution-guided library design (gray), showing how we densely sample the possible single and double mutations but only very sparsely sample higher-order mutants. (i) Histogram of the number of unique backgrounds on which each variant in the evolution-guided library is observed in the synthesized library based on sequencing analysis.

However, mutations distant from the binding site could also influence specificity, particularly in sequence backgrounds other than WT and M230A. Unfortunately, the space of combinatorial variants across the entire 436-residue DraNramp sequence is far too large to systematically explore. Instead, we leveraged information from natural sequences to prioritize specific changes. Based on existing work on generative protein sequence models^28,29^, we hypothesized that residue identities observed in natural sequences were more likely to preserve transporter function than those introduced through random mutagenesis. However, sampling directly from a range of state-of-the-art protein generative models decreased the chance of sampling the known specificity-altering mutations M230A/C/G/S/T compared to a random baseline (Fig. 1b-c). To address this, we developed an approach (latent space exploration) to sample the sequence space of natural transporters while biasing towards mutations more likely to alter function. Specifically, we built a simple site-independent fitness model that incorporates a bias toward mutations observed in distant, non-canonical Nramp homologs. We first constructed a latent space representation of a diverse multiple sequence alignment (MSA) of 17,763 Nramp homologs using principal component analysis (PCA; Fig. 1d; Extended Data Fig. 1a-d)^31^. Because the axes of this representation encode sets of mutations that capture the maximal variance in the alignment, we hypothesized that mutations causing large shifts in this space would be more likely to affect DraNramp function. The positions with the highest weights in this analysis are conserved within clades but not between clades (Extended Data Fig. 1e). When we incorporated into a generative sequence model framework an additional term to bias our sampling toward these positions (see *Methods*), the model then generated sequences with variation at the binding site and through the core of the protein (Fig. 1e), including a >10-fold enrichment for mutations at the known specificity-encoding position 230 (Fig. 1c).

While direct synthesis of a large library of sequences generated by this model was impractical, we used a variational synthesis approach^32^ to approximate sampling from this distribution. Using a stochastic library synthesis protocol, a limited set of mutations approximating the frequencies specified by the model were combined independently across 59 different sites of DraNramp (Fig. 1f). After bottlenecking, the final evolution-guided library contains 22,300 unique variants, and most variants have 1–5 changes relative to WT (Fig. 1g). This library densely samples the possible single and double mutations while randomly sampling the massive space of possible higher-order variants in the theoretical library design (Fig. 1h). The mutations at the designed sites plus additional random variation provided by synthesis errors yield a total of 1,797 distinct mutations, each observed on an average of 29 different backgrounds (Fig. 1i). Combining the site-specific library described above and this evolution-guided library, the entire combined library contains a total of 36,963 DraNramp variants.

### A riboswitch-based fluorescence assay allows for high-throughput screening of Mn^2+^ import

While several assays exist for Mn^2+^ import, none are optimized for high-throughput screening of large variant libraries. Therefore, we leveraged the regulatory mechanism of the *E. coli* Mn^2+^ exporter gene *mntP*^33^ to develop an *in vivo* fluorescence-based assay in *E. coli* that provides a robust and long-lasting signal, enabling cells to be sorted based on transporter activity. MntP protein levels are natively repressed both by the Mn^2+^-responsive transcription factor MntR^33^ and the riboswitch *yybP-ykoY*; high intracellular Mn^2+^ concentration relieves this repression, activating *mntP* expression^34^. We engineered a reporter plasmid, pSB1, by transplanting the *mntP* promoter and 5’ untranslated region containing the riboswitch upstream of the mEmerald coding sequence such that this green fluorescent protein is produced in response to high intracellular Mn^2+^ levels (Fig. 2a). pSB1-carrying cells overexpressing WT DraNramp, variants with intermediate Mn^2+^ transport (M230A and N59D), or negative controls (transport-inactive D56A variant or empty vector) showed a fast, medium, and slow Mn^2+^ concentration-dependent increase in green fluorescence, respectively (Extended Data Fig. 2a). These populations separate clearly by fluorescence measured by flow cytometry (Fig. 2b, Extended Data Fig. 2b), and we can estimate the effective Michaelis constant 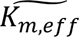 and relative 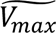 *in vivo* from the resulting dose-response curves (Fig. 2c). 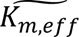 differs from the true microscopic Michaelis constant due to competition with other metals in the medium, especially Mg^2+^ and Ca^2+^, which are present at 1 and 0.25 mM, respectively. Nevertheless, values scale similarly to known *in vitro* kinetic parameters for DraNramp (Supplementary Table 1)^10^.

**Figure 2.**
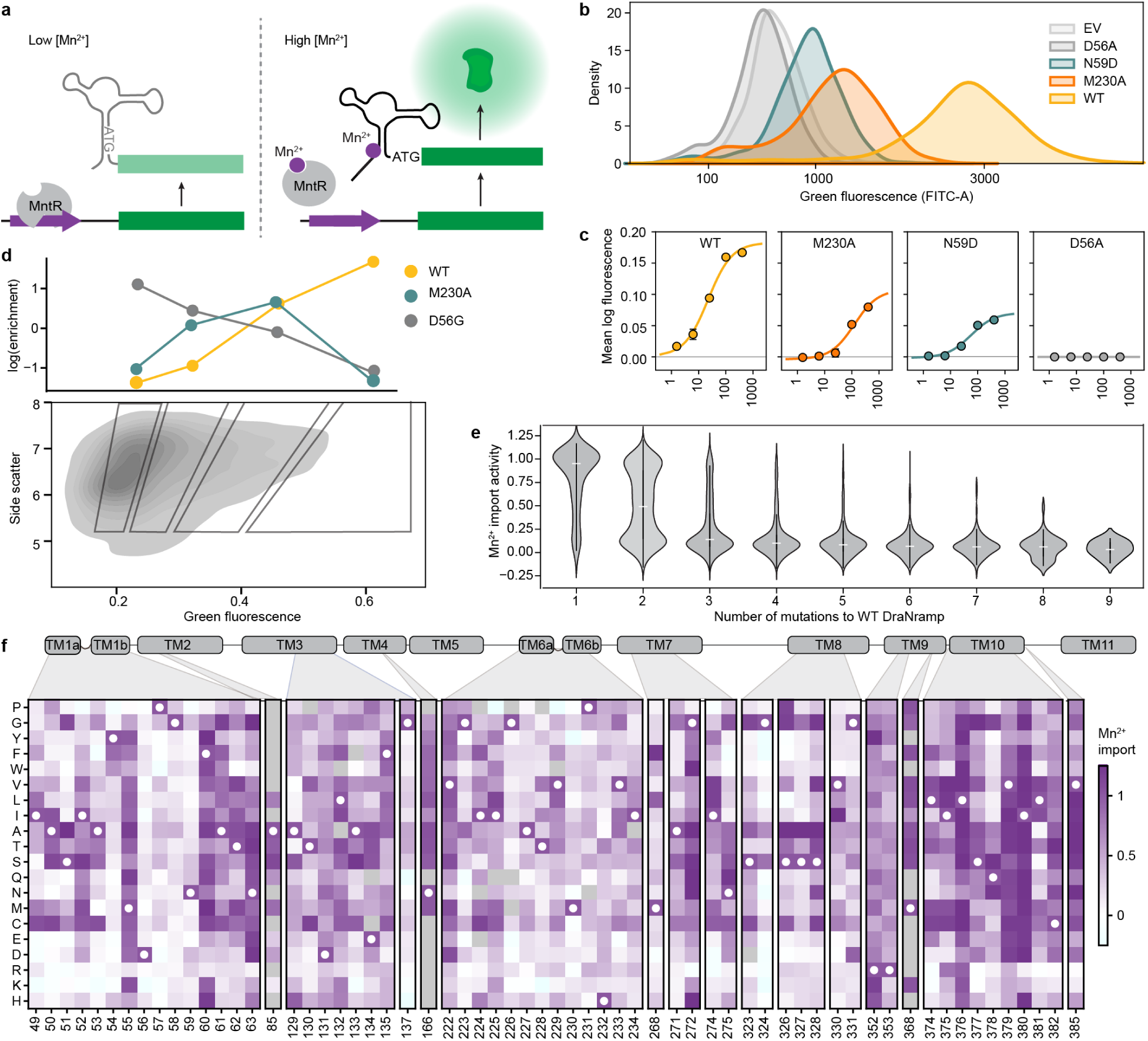
A high-throughput assay for Mn^2+^ import can map DraNramp’s local fitness landscape for its natural activity. (a) Schematic of our fluorescent reporter system that leverages both transcriptional and translational control to couple intracellular Mn^2+^ concentration to fluorescence. (b) Example green fluorescence distributions (kernel density estimates) measured by flow cytometry for *E. coli* cells expressing WT DraNramp (yellow), M230A (orange), and two variants with low (N59D; teal) or no (D56A; dark gray) transport activity, alongside the pET28a empty vector (light gray). The x-axis is plotted on a logicle scale and events were gated to have similar cell sizes as measured by side scatter. (c) Dose-response curves normalized to the D56A data relating MnCl_2_ concentration in the growth medium to fluorescence for the same variants as in (b), with overlaid fits to a sigmoid curve (resulting kinetic parameters listed in Supplementary Table 1). Error bars represent standard error of the mean from three replicates; sample raw distributions and an overview of the analysis are in Extended Data Fig. 2. (d) At the bottom is a kernel density plot showing the distribution of cells in the first replicate of the evolution-guided library screen on the correlated green fluorescence (FITC-A) and side scatter area axes; approximate locations of the four sorted bins are overlaid. Above is the log-transformed enrichment of WT and two variants with intermediate (M230A) or no (D56G) Mn^2+^ transport activity, highlighting how enrichment scores vary across bins for different variants. (e) Distribution of Mn^2+^ activity scores for substitutions in the evolution-guided library across mutational depth. Scores range between ∼0 to ∼1 (representing no activity to WT-like levels of activity), with scores above 1 representing improved activity and scores below zero likely representing experimental noise. (f) Heatmap of Mn^2+^ import scores for all variants with single mutation at positions with at least 5 measured variants, primarily from the binding-site library. White circles mark wildtype amino acids. Gray positions lack data.

We screened our two DraNramp variant libraries through our Mn^2+^ import assay. We sorted cells grown in the presence of 100 µM MnCl_2_ into four bins representing low, intermediate, wildtype, and increased fluorescence levels. Sorting thresholds were based on control fluorescence distributions for cells expressing WT DraNramp, the inactive control D56A, and the moderate-activity variant N59D (Fig. 2d; Extended Data Fig. 3a-b). We sequenced these sorted populations to calculate a relative Mn^2+^ import score in which WT DraNramp has a score of 1 and inactive controls a score of 0 (Extended Data Fig. 3c-d). These scores distribute broadly (Extended Data Fig. 3e-f), correlate well between replicates (Extended Data Fig. 3g-h; R^2^ = 0.61-0.90) and qualitatively recapitulate the effects of known variants tested *in vitro* and in other *in vivo* assays (Supplementary Table 1), suggesting that they provide a robust and accurate readout of Mn^2+^ import activity across the entire library. Variants present in both libraries also show correlated scores (Extended Data Fig. 3i), allowing us to merge all the data into a single dataset containing the Mn^2+^ import of 36,963 DraNramp variants.

As expected, the distribution of mutational scores in the evolution-guided library drops off with mutational depth, with most variants with three or more mutations being inactive (Fig. 2e). Single-mutation variants in the evolution-guided library transport more Mn^2+^ on average than those in the binding-site library (Extended Data Fig. 3e-f). Among these binding-site mutational changes, the conserved binding-site residues (residues 56-59 in TM1 and 230-232 in TM6) are highly constrained, with notable exceptions (Fig. 2f). While aspartate is strictly required at position 56 for Mn^2+^ transport, substitutions of asparagine at position 59 with other polar residues or even glycine, but not glutamate, still allowed for Mn^2+^ transport. Additionally, most variants with substitutions in TM10a (374-385) have near-wildtype or even increased Mn^2+^ import levels (Fig. 2f and Extended Data Fig. 4). This pattern is intriguing because TM10a moves independently in the conformational change between outward- and inward-open^6,8,35^. Substitutions at Q378—which forms a water-mediated contact with Mn^2+^ ^8^—and L381 are exceptions to this TM10a pattern, with many deleterious substitutions. Of note, DraNramp L381 corresponds to a glutamine in human Nramps that directly coordinates Mn^2+^ in recent structures^5^. Overall, the observed patterns of single-mutation effects show that the assay successfully recapitulates expected patterns. The patterns also paint a general portrait of residues closest to the metal-binding site contributing most to the protein’s native activity (Extended Data Fig. 4c), transport of its physiological substrate Mn^2+^, and hint that effects on the conformational equilibrium may also impact metal ion transport.

### For most variants, Mn^2+^ import activity can be explained by additive effects of single mutations

Many individual mutations in the library are observed on multiple backgrounds, allowing us to assess their effect on Mn^2+^ import activity in isolation and how those activities change when mutations are combined. A simple extrapolation of single-mutation effects to higher-order mutants poorly predicts their activity (Extended Data Fig. 5a-c). We therefore used a regression framework to estimate background-averaged mutational effects. A linear regression model improves over the simple extrapolation but still fits poorly (Fig. 3a) as this model does not account for global epistasis inherent in the likely nonlinear genotype-to-phenotype relationship in our dataset. Many studies have addressed global epistasis by assuming that mutation effects combine additively with respect to an unobserved latent activity (or “fitness potential”^36^), which is then nonlinearly related to the observed phenotype^36-39^. While this nonlinear function can take many forms, we chose a sigmoid for its simplicity and simple thermodynamic interpretation. To estimate posterior distributions robustly, we fit this model using Bayesian regression via stochastic variational inference incorporating experimental error and assigning normal priors to the regression parameters (see Supplementary Note 1 for a detailed explanation of our inference approach).

**Figure 3.**
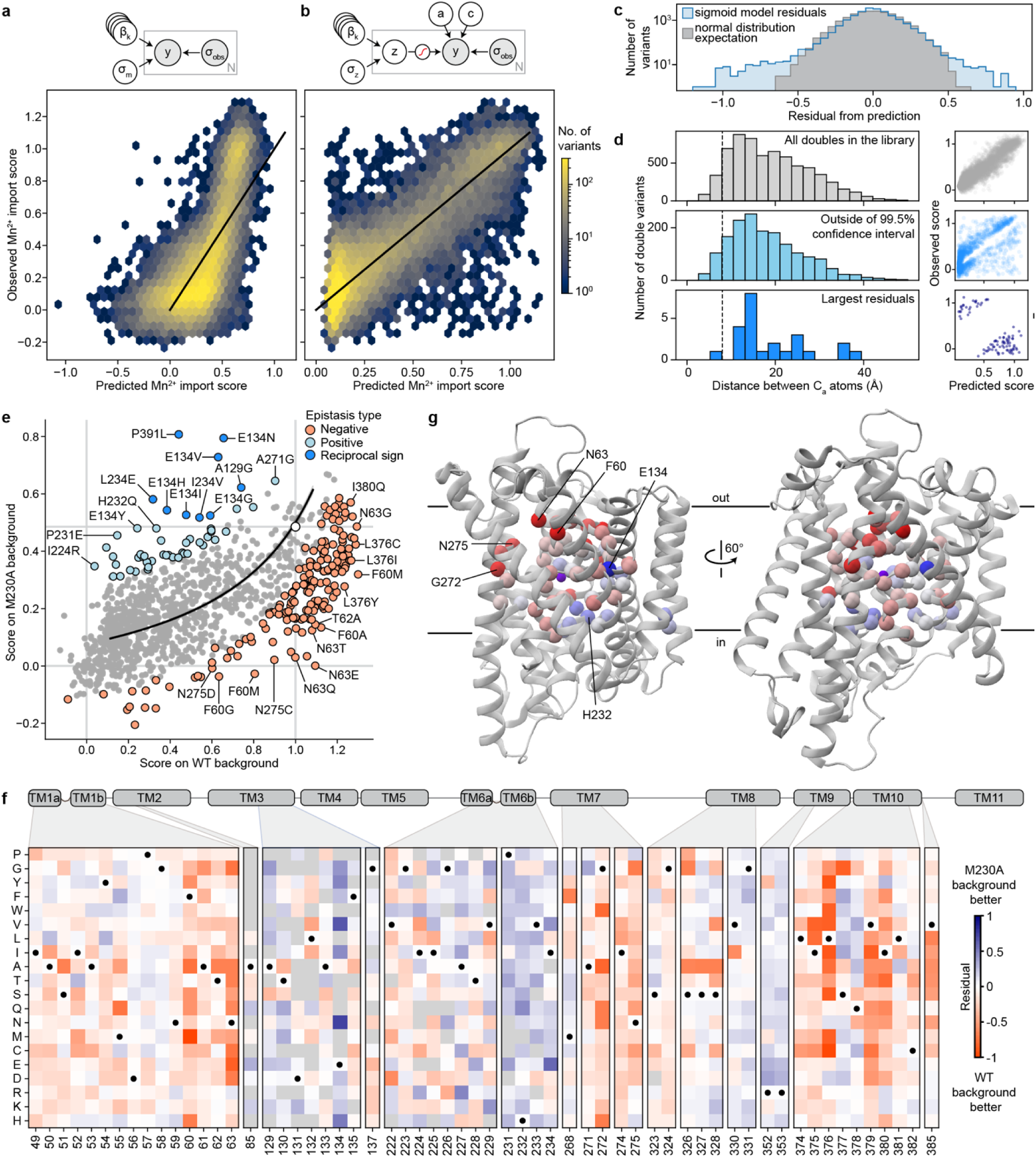
Modeling the DraNramp mutational landscape for Mn^2+^ import. (a-b) Density plots comparing predicted and observed Mn^2+^ import scores for variants from the combined library when modeled with linear regression (a) and the sigmoid regression model (b). A black line indicates perfect predictions. A graphical model of the Bayesian regression approach is shown above each graph, using plate notation to indicate the different data points (See Supplementary Note 1 for a detailed description). Predictions were made as part of a randomly held-out set of 20% of the data, and points with estimated experimental error values greater than 0.15 are hidden. (c) Distribution of the sigmoid model residuals (blue) compared to expectation from normally distributed errors with the same mean (gray). (d) Distributions of C_α_-C_α_ distances based on the occluded DraNramp structure (PDB ID: 8E60) for double mutants in the combined library (top, light gray), double mutants with residuals outside of the 99.5% confidence interval predicted from their experimental error and the global epistasis model posteriors (light blue), and double mutants with absolute residuals >0.5 (darker blue). Dashed vertical lines indicate a sample 8-Å cutoff for structural contacts, and on the right is the locations on the plot in (b) of the points represented by each histogram. Mutation pairs with stronger epistasis are not structurally closer on average. (e) Scatterplot showing the score of single mutants on the WT (x-axis) versus the M230A (y-axis) background. A black line represents the fit of the sigmoid global epistasis model. Samples within a 95% confidence interval of the model’s prediction are shown in gray but are nearly invisible due to the confidence interval’s small size. Mutations highlighted in light red exhibit negative epistasis with a residual of <0.2 below the model prediction, while those in light blue have residuals of >0.2 above the model prediction. Those in darker blue have a reciprocal sign epistasis relationship with M230A. The point representing WT and M230A on their own is shown in white, and the WT and M230A activity levels are shown as light gray lines. (f) Heatmap of residuals for all double mutants with M230A in the combined library at positions with a Mn^2+^ import score for at least seven different amino acid changes on the WT and M230A backgrounds. Black circles mark wildtype amino acids. Gray positions lack data. (g) The average residual at each position is mapped onto the DraNramp structure. The color scales for (f) and (g) are the same.

While adding only two parameters, the sigmoid model dramatically improves model fit, approaching the limit set by experimental noise (R^2^=0.73, Fig. 3b and Extended Data Fig. 5b). Interestingly, some observed variants deviate from the model’s predictions more than accounted for by normally distributed errors (Fig. 3c), suggesting that these variants represent cases of significant specific epistasis.

### Outliers representing specific epistasis cluster on the DraNramp structure

To investigate the outliers of our global epistasis model, we analyzed all 8,236 double-mutant cycles in the dataset. Approximately one in four double-mutant cycles showed significant epistasis (outside of a 99.5% posterior predictive interval), and these epistatic double-mutant cycles do not have a substantially different inter-residue distance distribution than all pairs (Fig. 3d). This is also true for the 250 mutation pairs exhibiting reciprocal sign epistasis, which are distributed throughout the protein (Extended Data Fig. 5d). Using pairwise couplings terms inferred from the double-mutant cycles to predict the outcomes of the 1,231 triple-mutant cycles in the dataset, we again find many large three-way epistatic residuals (Extended Data Fig. 5e-g), indicating that pairwise epistasis is insufficient to describe the observed variation in the triple-mutant dataset.

To gain more insight into potential mechanisms underlying this epistasis, we focused on double-mutant cycles including M230A, which are abundant in our binding-site library, including most residues in the first and second shell around the binding site (Extended Data Fig. 4). The sigmoid epistasis model generally explains the effects of mutations on the WT and M230A backgrounds, but with many outliers representing both positive and negative epistasis (Fig. 3e). Nine mutations—including four at position 134—show a reciprocal sign epistasis relationship with M230A. We calculated a residual for each variant as the difference between its predicted and observed score on the M230A background (Fig. 3f). Mutations to TM1b, TM7, and one face of TM10a tend to transport Mn^2+^ less well than expected on the M230A background, while mutations to TM2 (especially residue 134), TM6b, a different face of TM10a, R352 and R353 tend to transport better than expected. Notably, E134, R352, and R353 are all important for proton cotransport^10^, while TM6b is important for metal and proton release and mutations at several of its residues bias DraNramp toward an inward-open conformation^40^. Plotting the average residual by position on the occluded structure of DraNramp (PDB ID: 8E60), the residues exhibiting negative epistasis tend to cluster around the outer vestibule (Fig. 3g), suggesting that mutations at these residues may preferentially destabilize conformations in which the outer vestibule is closed. This trend and the opposing positive epistasis observed at inward-stabilizing TM6b positions suggest a possible role for DraNramp’s conformational equilibrium between its inward- and outward-open state in shaping the observed specific epistatic trends.

### A growth complementation assay allows for high-throughput screening of Mg^2+^ import

Our experimental map of Mn^2+^ import in DraNramp helps us understand what features control import of one of DraNramp’s native substrates. However, it does not explain how DraNramp excludes Mg^2+^ or what kinds of mutations would alter this specificity. To do so, we screened our libraries of DraNramp variants for Mg^2+^ import activity. We were unable to adapt the riboswitch-based assay to intracellular Mg^2+^ sensing despite the existence of Mg^2+^ riboswitches^41^. Reliably measuring Mg^2+^ import by DraNramp variants would require competition with *E. coli*’s far more efficient existing Mg^2+^ import machinery, against a background of much higher intracellular ion concentrations of Mg^2+^ than Mn^2+^ ^42,43^. We instead adapted a Mg^2+^ complementation assay originally developed to assess the activity of the Mg^2+^ transporter MgtE^44^. In this system, transporter variants are expressed in a Mg^2+^-auxotrophic *E. coli* strain lacking all three of its native Mg^2+^ importers^44^ (BW25113 Δ*mgtA* Δ*corA* Δ*yhiD*, in which we deleted *recA* to generate a strain henceforth referred to as MgKO). In the absence of an additional transporter, these cells require 100 mM MgSO_4_ to survive, but they can grow in 1 mM MgSO_4_ if expressing a functional transporter (Fig. 4a-b).

**Figure 4.**
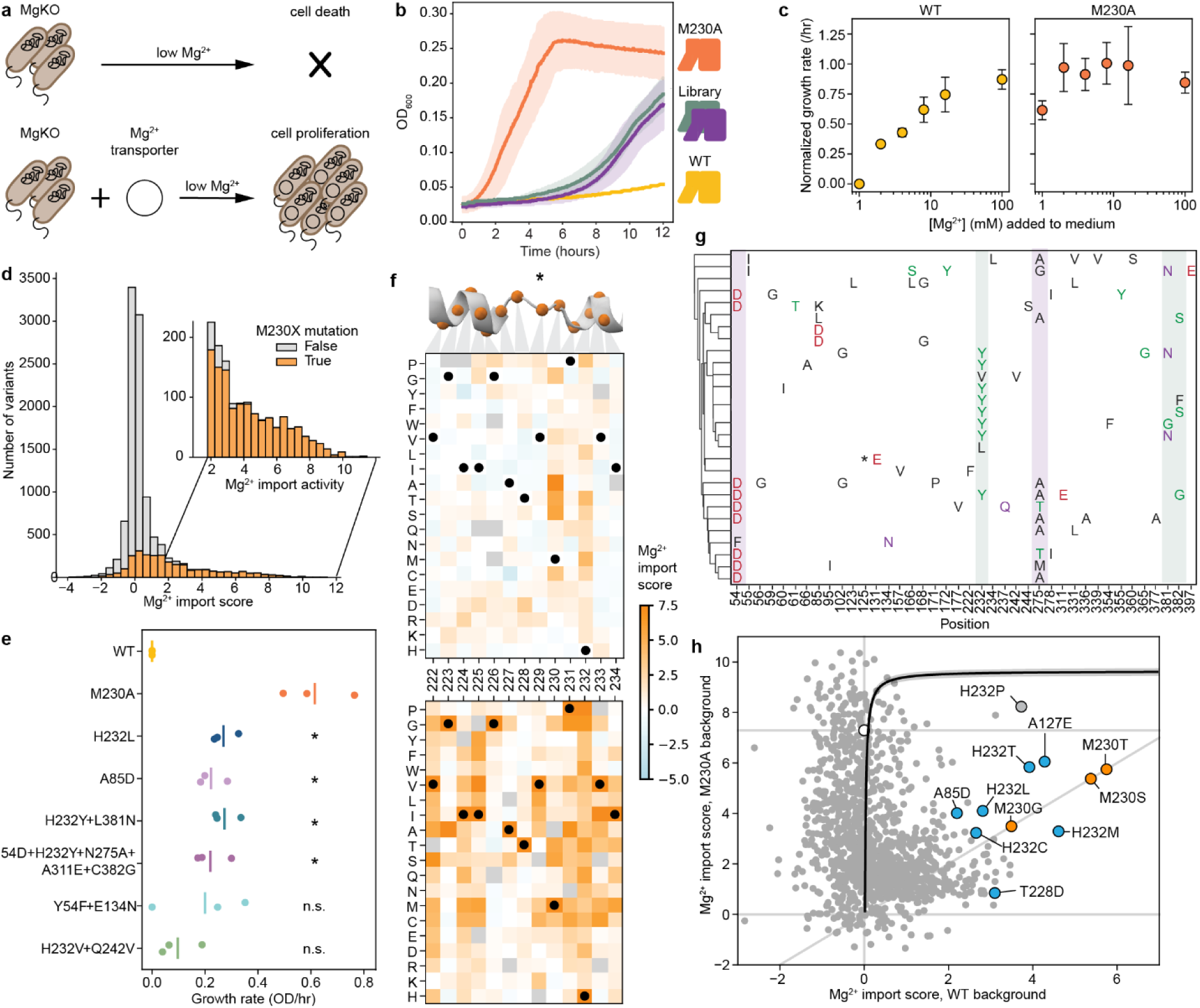
A growth complementation assay facilitates mapping of mutations promoting Mg^2+^ import in DraNramp. (a) MgKO, a Mg^2+^-auxotrophic strain of *E. coli* lacking any genetically encoded Mg^2+^ transporters, can only survive in low Mg^2+^ when rescued with a functional Mg^2+^ transporter. (b) Example growth curves in LB supplemented with 1 mM MgSO_4_ with 20 µM IPTG, showing robust growth for the Mg^2+^-transporting M230A variant (orange), no growth for the non-transporting WT DraNramp (gold), and intermediate growth for two replicates of the evolution-guided library (purple and teal). (c) Growth rates, normalized to WT in 1 mM supplemental MgSO_4_, measured across concentrations of Mg^2+^ supplemented into LB medium. Rates are higher for M230A (right) compared to WT (left). Error bars represent standard error of the mean across three replicates. (d) Distribution of Mg^2+^ import scores from the combined libraries, colored by whether the M230 position is mutated, with the region with scores above 2 (representing clear Mg^2+^ import) in an inset. (e) Growth rates of isolated clones for a selected set of variants with no M230 mutation that have Mg^2+^ import scores significantly higher than WT. M230A is included as a positive control. Stars represent significance thresholds from a series of Welch’s t-tests against the WT with a Benjamini-Hochberg correction (*: p<0.05). (f) Heatmaps of Mg^2+^ import scores for all mutations to TM6 on the WT (top) and M230A (bottom) backgrounds. Black circles mark wildtype amino acids. Gray positions lack data. The 230 column is identical between heatmaps. Above the heatmaps is a snapshot of TM6 (PDB ID: 8E6N). The black asterisk approximates the bound Mn^2+^; Mg^2+^ may or may not bind the same site in the M230A variant. (g) Twenty-eight variants from the evolution-guided library with scores significantly outside the distribution defined by WT barcodes (FDR < 0.05 via Student’s t-test with a Benjamini-Hochberg correction). Variants were clustered based on which residues (color-coded by chemical property) were mutated using clustermap in seaborn. Arrows above and corresponding boxed columns highlight the positions of observed sequence couplings (purple: positions 54 and 275; green: positions 232 and 381/382). The asterisk represents a deletion of residues 125-127. (h) Scatterplot of Mg^2+^ import scores for all single mutations present on both the WT (horizontal axis) and M230A (vertical axis) background. The gray diagonal line represents mutations with the same overall activity level on both backgrounds, while the horizontal gray lines represent the WT score (0.00 ± 0.04) or M230A score (7.29 ± 0.62). The black curve represents a fit to a site-independent sigmoid model, demonstrating how the mutations deviate considerably from an additive model even accounting for sigmoidal global epistasis. The shaded region around the sigmoidal curve represents a 99.7% confidence interval. A selection of mutations that improve Mg^2+^ on their own are highlighted in teal, and several M230 mutations (which by definition have the same effect on each background) in orange.

M230A-expressing cells grow better than WT-expressing cells across a wide range of Mg^2+^ concentrations, with 1 mM being the lowest concentration at which WT-expressing cells consistently do not grow (Fig. 4c). We transformed the binding-site and evolution-guided libraries into MgKO cells and sequenced plasmids extracted from cells after twelve hours of growth in 1 mM MgSO_4_. We calculated log enrichment as Mg^2+^ import scores, comparing variant abundance pre- and post-selection.

We observed large stochastic variation with many variants (50-80%) dropping out of each replicate in the pre-selection overnight growth, likely due to the compromised viability of MgKO cells, although we still have at least two observations for 10,714 variants. However, more than half of TM3 and TM6 variants dropped out in the initial binding-site library screens, so we repeated the screens on these isolated subpools and increased coverage of the variants in these subpools from 49% to 93%. Replicates screens showed good correlation (Extended Data Fig. 6a-e), and in all replicates the positive control variants M230A, M230S, M230T, and M230G scored higher than WT, with average enrichments of 31-fold and 1488-fold in the binding-site and evolution-guided libraries, respectively. This variation in enrichment scale is due to the larger fraction of Mg^2+^-transporting variants in the binding-site library, which contains a higher proportion of M230A double mutants. As with the Mn^2+^ dataset, we used linear regression to rescale and merge the results from these two libraries into a combined dataset for overall comparison, and report both the original and combined scores in Supplementary Table 2.

In the combined dataset, most variants with very high Mg^2+^ import scores contain mutations at position 230 (Fig. 4d). Despite this dominance of M230X mutations, 37 variants without M230 mutations were significantly enriched for improved growth in Mg^2+^-limiting conditions (Supplementary Table 3), using the individual barcodes scores corresponding to WT as a non-transporting null distribution (Extended Data Fig. 6f-h). We cloned and retested several of these variants, and all had improved growth relative to WT at 1 mM MgSO_4_ (Fig. 4e, Extended Data Fig. 7a), and across a range of Mg^2+^ concentrations (Extended Data Fig. 7b). This validates that even relatively small but statistically significant enrichment represents improved growth, although it does not, on its own, show that this growth is due to Mg^2+^ import. However, we do observe that the growth improvement decays as the stringency of the Mg^2+^ selection decreases both from screening data (Extended Data Fig. 7c) and for the subset of variants tested individually (Extended Data Fig. 7d), suggesting that the growth improvement is tied to Mg^2+^ and not due to a general increase in fitness of the cells.

Among single variants, M230A/S/T have the highest scores, followed by mutations to H232 (Fig. 4f and Extended Data Fig. 8a-b). While the role of M230 in specificity was already known^1^, the role of H232 has been less clear, even though this histidine is highly conserved in canonical Nramps (Extended Data Fig. 1e). H232 lies below the metal-binding site and is not known to directly coordinate the ion but has been suggested to act as a pivot point in conformational change^40^ and as a proton-binding site^16^. A striking diversity of mutations at this position enables MgKO growth, including mutations to small polar amino acids (S, T), but also to P, M, Q and N. H232M, introducing a second methionine in the TM6 metal-binding region, is especially surprising. Interestingly, a spontaneous variant with an alanine inserted before M230 (ins230A) also rescues MgKO growth despite still retaining M230. Together, these clusters of high-scoring variants at positions 230 and 232 suggest that the mechanisms by which DraNramp excludes Mg^2+^ are not as simple as increasing the binding-site size or removing a sulfur atom. Many other mutations have the potential to change the binding-site size and preserve Mn^2+^ import. These results instead suggest that the particular placement of a methionine and histidine together create a binding site that efficiently transports Mn^2+^ and excludes Mg^2+^, and several ways of breaking this arrangement can allow for Mg^2+^ import.

While all significant hits from the binding-site library were in TM6b, particularly at positions 230 and 232 (Fig. 4d and Extended Data Fig. 6e-f), additional variants from the combinatorial library also scored high enough to suggest they rescued growth, including 2 we isolated and confirmed had growth rates higher than WT but below M230A, as expected from the screen results (A85D and Y54F+E134N; Fig. 4e). The M230-retaining variants with significant Mg^2+^ import scores include one single variant (A85D) and 27 combinatorial variants from the evolution-guided library (Fig. 4g). Many of these 28 variants contain mutations at Y54, A85, H232, N275, L381, or C382. Notably, six of eight Mg^2+^-importing variants containing Y54D also have mutations to N275, and four of five containing H232Y also have mutations at either L381 or C382 (Fig. 4g). These co-occurrences suggest energetic couplings within each of these residue pairs. In both cases the pairs are structurally close without direct sidechain contact, although N275 hydrogen-bonds with the Y54 backbone (Extended Data Fig. 8c-d). This suggests that some specificity-switching mechanisms require multiple mutations, although why mutations at these particular pairs are so successful is not immediately clear.

To begin to better understand the landscape of Mg^2+^ import in DraNramp, we compared the effect of single mutations on the WT and M230A backgrounds (Fig. 4f and 4h). Starting from the Mg^2+^ import-competent M230A background, most mutations either have no effect or decrease the Mg^2+^ import score, with few increasing it. The most deleterious mutations cluster at the binding site, particularly at positions 226-228 in the unwound region of TM6 (Fig. 4f). The A227 backbone carbonyl coordinates the Mn^2+^ ion^8^, and our results suggest that Mg^2+^ import on the M230A background is particularly sensitive to A227 positioning. Conversely, positions 231-232 (the “PH” in the highly conserved MPH motif) become highly permissive to mutations on the M230A background (Fig. 4f). That different residues in the unwound region of TM6 are essential for Mg^2+^ and Mn^2+^ import suggests that this region mediates different binding modes for the two ions, with Mn^2+^ more reliant on the MPH motif and Mg^2+^ on the positioning of the upstream residues.

Additionally, most mutations affect the Mg^2+^ import score differently on the WT and M230A backgrounds (Fig. 4h). This contrasts with the Mn^2+^ import results where mutations tend to combine additively (Fig. 3b and 3e). Mutations like H232M/T/S/L and A85D show negative sign epistasis, with similar Mg^2+^ import scores on both the WT and the M230A backgrounds: adding the M230A mutation to these H232M/T/S/L and A85D variants has little to no effect, but adding the corresponding mutations to the M230A background is deleterious (teal datapoints in Fig. 4h; Extended Data Fig. 8e). To better understand the mechanisms that give rise to the non-additive complexity of these mutational effects, we turned to comparing the datasets with the two different substrates, Mn^2+^ and Mg^2+^.

### Disentangling determinants of global transport function and substrate specificity

We sought to differentiate mutations that affect transporter function globally—for example, via folding and overall conformational cycling—from ones that control substrate specificity. Mg^2+^-transporting variants showed a wide range of Mn^2+^ transport activity (Fig. 5a). Most M230X-containing variants (orange) follow a general trend where their Mn^2+^ and Mg^2+^ activity scores range from ∼0—representing transporters inactive in our assays—to the most active M230A/S/T variants. This trend suggests that mutations to M230 primarily alter specificity, whereas mutations elsewhere often affect folding or other determinants of global function. However, some mutations, when combined with M230X, deviate from this trend, including the handful of Mg^2+^-transporting M230-retaining variants discussed in the previous section.

**Figure 5.**
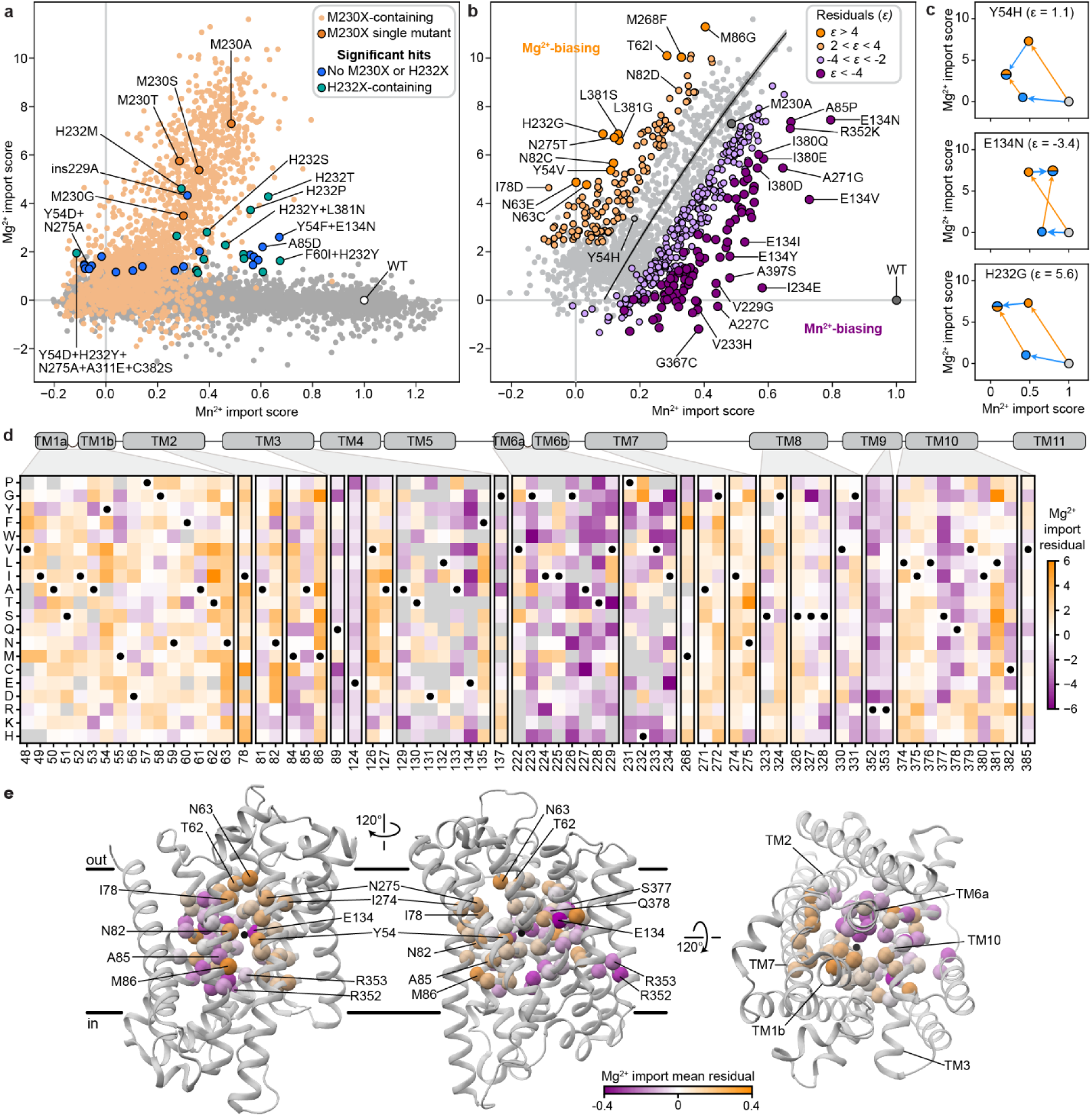
Disentangling global and substrate-specific contributions to activity. (a) Comparison of the Mn^2+^ versus Mg^2+^ import scores for all variants in the combined libraries. Variants containing mutations to residue 230 are highlighted in orange, with most other variants in gray. In blue (no H232 mutation) or teal (with M232X mutation) are a subset of the highest-scoring M230-retaining variants with statistically significant Mg^2+^ import (FDR < 0.05). (b) Scatterplot of the Mn^2+^ versus Mg^2+^ import scores for all double mutants containing M230A. The black line represents fit of the sigmoid specificity model and points are colored according to residuals from this null expectation. (c) Double-mutant cycles in specificity space for three example mutations combined with M230A. In each case, orange arrows represent the M230A mutation while blue arrows represent the highlighted mutation. Y54H has a small residual from the expected specificity relationship (*ϵ* = 1.1); E134N has a very negative residual (Mn^2+^-biasing, *ϵ* = −3.4); and H232G has a very positive residual (Mg^2+^-biasing, *ϵ* = 5.6). (d) Heatmap of residuals as calculated in (b). Darker orange represents greater Mg^2+^ import than expected and darker purple represents greater Mn^2+^ import than expected. Black circles mark wildtype amino acids. Gray positions lack data. (e) Average residual scores from (d) mapped onto the occluded structure (PDB ID: 8E60). Positions and TMs discussed in the main text are labeled.

To determine which residues contribute to specificity, we analyzed the effects of M230A-containing double mutants on Mn^2+^ and Mg^2+^ transport. We adapted our sigmoid global epistasis model to account for multiple substrates by assuming equivalent latent functional effects but a differentially scaled sigmoid function for each substrate (see Methods and Supplementary Note 1). We could then fit additional specificity parameters for individual variants. For instance, we fit an overall curve for the expected relationship between all M230A double mutants assuming that only M230A perturbs the latent specificity (Fig. 5b, black line). While many mutants follow this trend, others deviate in a manner that suggests either Mg^2+^- or Mn^2+^-biasing.

As with our previous epistasis analysis, we calculated residuals from this initial expectation (Fig. 5b-e). At certain positions, most or all mutations have similar effects on specificity in the M230A background. Positions in TM6 upstream of M230 are essential for Mg^2+^ but not Mn^2+^ import and thus mutations at those positions drive the M230A variant toward Mn^2+^ specificity. Mutations to many proton-pathway residues (E134, R352, R353) or TM10 (S377 and Q378) also enhance Mn^2+^ preference, whereas substitutions at T62 (TM1b) above the metal-binding site, M86 (TM2), N275 (TM7)—part of the outer gate—and L381 (TM10) generally strongly push M230A toward Mg^2+^ specificity. Some mutations—especially those introducing more polar or negatively charged amino acids (Y54E/C/M/N/Q, I78D/H, N82E/Q, A85D/E, M86Q, I274N, and L381Q/N/M/G)—allow Mg^2+^ transport without measurable Mn^2+^ import. Except for A85D, these mutations do not increase Mg^2+^ import on their own, but they increase Mg^2+^ specificity on the M230A background (Fig. 5b, Extended Data Fig. 8a-b)). This is consistent with the idea that a major biochemical perturbation near the metal-binding site is required for M230X-like Mg^2+^ import activity, after which secondary mutations can more easily modulate specificity.

### Specificity-altering mutations broadly fall into two classes

The discrepancy between the small set of core mutations that break DraNramp’s selectivity against Mg^2+^ and the large number of positions that modulate specificity on the more promiscuous M230A background suggests that specificity determinants in DraNramp fall into multiple classes. To gain a data-driven perspective on the core specificity determinants, we modified the sigmoid specificity model so it is no longer constrained to only fit terms at position 230. Briefly, this specificity regression model fits a set of global regression parameters *β*_*g*_ that attempt to explain both the Mn^2+^ and Mg^2+^ datasets, while a second specificity set, *β*_*s*_, fits only the most essential terms to explain the difference between the two datasets (Fig. 6a; Extended Data Fig. 9). We set a sparse Laplace prior on the *β*_*s*_ parameters to encourage the model to infer as much of the variance as possible on a sparse set of specificity determinants.

**Figure 6.**
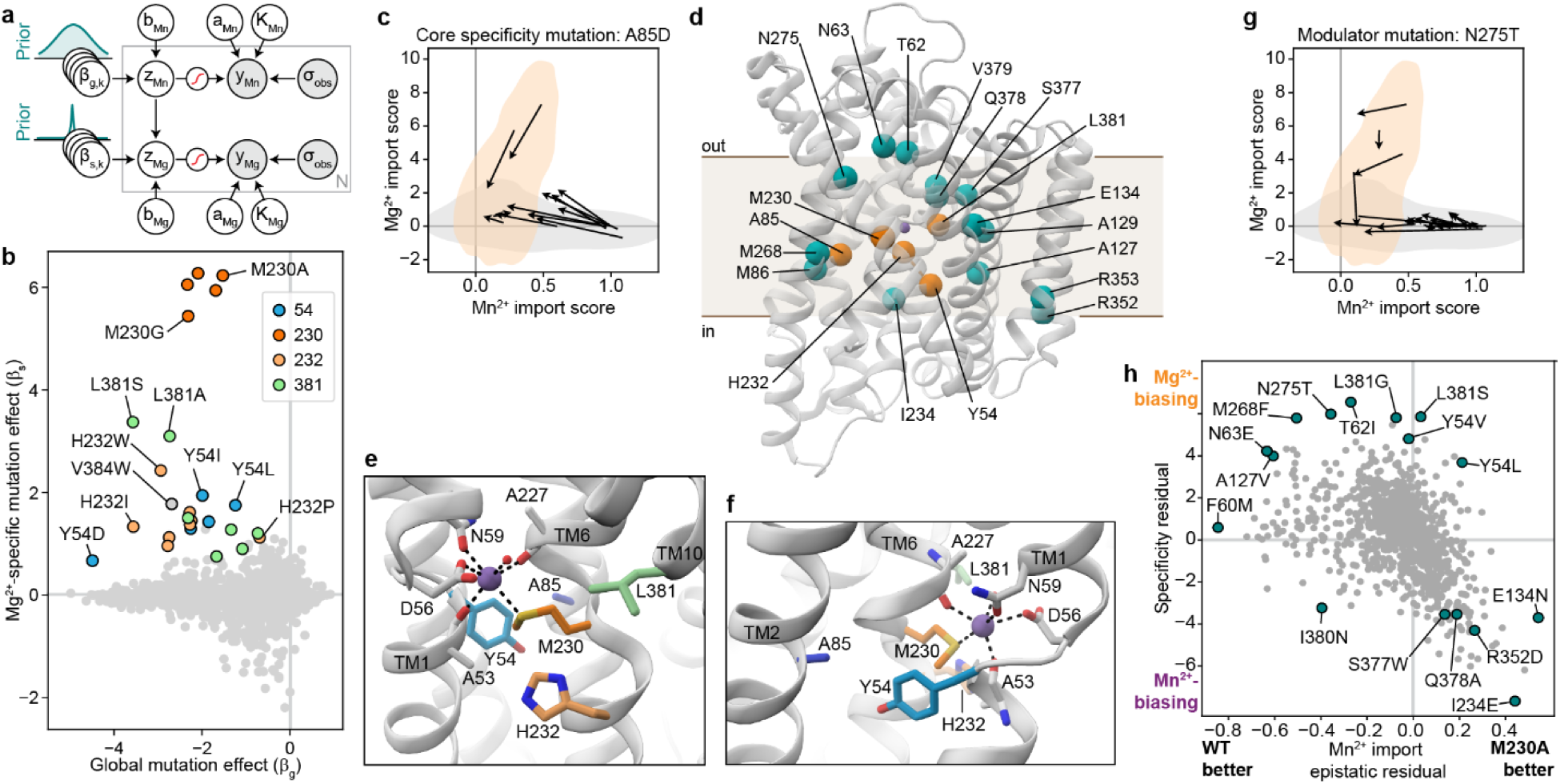
Identifying core and modulator specificity positions in DraNramp. (a) Schematic of specificity regression model, highlighting two sets of regression parameters *β*_*g*_ and *β*_*s*_. The different priors on these two sets of parameters (Gaussian and Laplace) are highlighted behind them. Y-axes are not to scale. (b) The learned weights of the global mutation effect terms (*β*_*g*_) fit to both Mn^2+^ and Mg^2+^ versus the sparse Mg^2+^-specific mutation effect terms (*β*_*s*_). Regression coefficients for positions 54, 230, 232, or 381 are highlighted; all other mutations are in gray. (c) Arrow field plot for A85D, in which each arrow defines a pair of variants differing by the A85D mutation and the arrow represents the effect of adding A85D on both Mn^2+^ and Mg^2+^ import. Behind the arrows, density contour lines for variants without M230 mutations (gray) or with M230 mutations (orange) are highlighted for comparison to Figure 5a. (d) The occluded structure of DraNramp (PDB ID: 8E60) with the five “core” specificity positions as orange spheres and notable specificity modulator positions in teal. (e-f) The core positions and additional key metal binding residues D56 and N59 on the same structure, colored as in (b). (g) Arrow field plot for a modulator specificity mutation, N275T, showing how the N275T mutation has a variety of effects both epistatically when considering a single substrate and specifically when considering both. (h) Scatterplot comparing the epistasis residuals from Figure 3g with the specificity residuals from Figure 5d; each point represents a single mutation. The two variables correlate with Pearson’s R = -0.44.

The model identifies four Mg^2+^ selectivity determinant positions: Y54, M230, H232, and L381 (Fig. 6b, Extended Data Fig. 9f). Mutations at each of these positions allow for Mg^2+^ import on their own or when combined with one other mutation (Y54D+N275A and Y54F+E134N; M230A/S/T/G/C; H232T/S/P/M/L/I; and L381N+H232Y). Mutations at these positions also increase the Mg^2+^ specificity of the Mg^2+^-transporting M230A variant (Fig. 5b). However, the model fails to identify some mutations because it assumes that the latent activity and specificity effects of each mutation are independent of mutational background. A salient example is A85D, which increases DraNramp’s specificity toward Mg^2+^ across many different backgrounds, but not on M230A or M230G (Fig. 6c). We nonetheless group position A85 with Y54, M230, H232, and L381 as the set of “core” specificity residues (Fig. 6d-f).

Mutations at these core positions can disrupt how DraNramp natively selects against Mg^2+^ import. The identities of these mutations are not consistent with Mg^2+^ simply requiring a larger binding site to accommodate a more hydrated ion. While M230A likely enlarges the binding site, other mutations allow for Mg^2+^ import without enlarging the binding site, and many mutations that likely enlarge the binding site and still transport some Mn^2+^ (such as N59G or V228A) do not lead to any detectable Mg^2+^ transport (Extended Data Fig. 8b). Outside of TM6, mutations at core position Y54 may directly interact with the metal or reposition the key metal-coordinating residue D56. Core positions A85 and L381 both sit in the second shell with the potential to interact more closely with the metal at some point in the conformational cycle. Indeed, L381 is an asparagine in both human Nramps and was found to directly coordinate the metal in a recent structure of human NRAMP2 ^5^. However, the fact that both L381N and L381G affect specificity suggests that their effects are unlikely to depend on a particular chemical interaction.

Many other mutations have highly context-dependent effects on specificity, unable to independently disrupt WT DraNramp’s exclusion of Mg^2+^ but able to finetune specificity once that threshold has been crossed. For example, N275T does not allow for Mg^2+^ import unless combined with core mutations M230X or Y54D, and on an M230X background sometimes, but not always, increases Mg^2+^ specificity (Fig. 6g). We refer to these as “modulator” positions; they are widely spread throughout the protein (Fig. 6d). What remains unclear from this analysis is how such mutations distant from the binding site can affect specificity.

A clue to the effect of these modulator positions lies in their overlap with key epistatic positions identified above (Fig. 3b). We compared the Mn^2+^ import epistatic residuals (Fig. 3f) of M230A double mutants with their specificity residual (Fig. 5d). These two independent metrics are significantly negatively correlated (Fig. 6h), meaning that residues that import Mn^2+^ better than expected in the M230A background tend to import Mg^2+^ less well than expected in the M230A background, and vice versa. This suggests that the same causative factor could underly the observed complex structure of the epistatic landscape and the specificity landscape.

### A core conformational balance mechanism can underly epistasis and specificity modulation

Based on our observations that key epistatic and specificity positions overlap, we hypothesized that mutations at these positions could cause differential conformational biasing from Mn^2+^ and Mg^2+^. We considered mathematically how variation of the energy difference between the inward- and outward-open states, Δ*G*_*flip*_, would affect the V_max_ of a transporter given a simple four-state model of its reaction cycle (Fig. 7a). We made three assumptions: (1) transporter conformational change is rate-limiting relative to substrate binding and release, (2) all intermediates are at steady state, and (3) energetic differences to the conformational flip transition state are linearly related to the energy difference between the outward- and inward-open conformations by a constant value 0 < *ϕ* < 1^45^ (Fig. 7b). We thus arrived at Equation 1 (see full derivation in Supplementary Note 2):

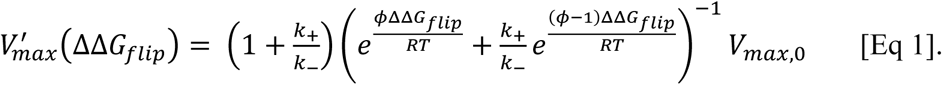

**Figure 7.**
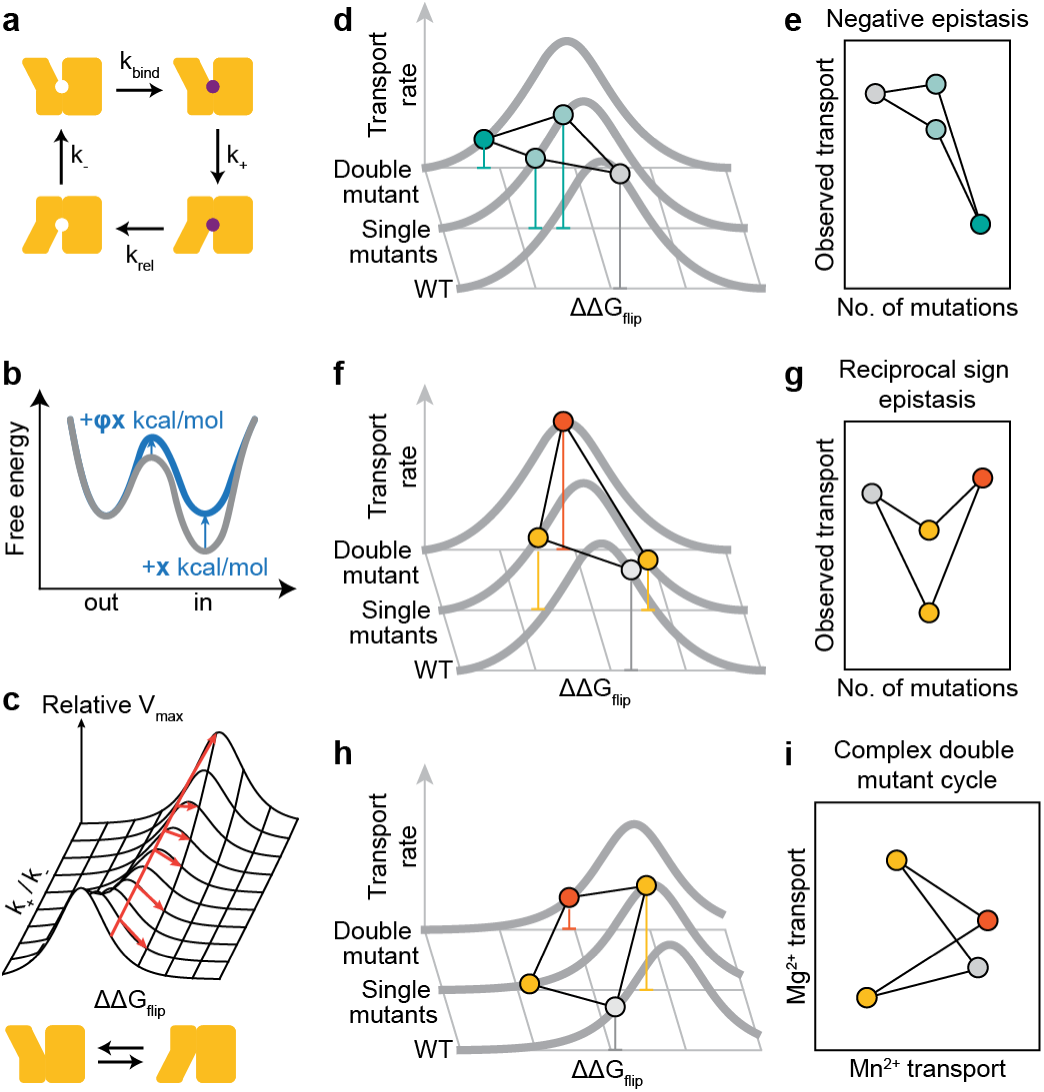
Transporter conformational balance can modulate specificity. (a) Schematic of transport model used for the conformational balance equation. (b) Reaction scheme with (blue) and without (gray) a mutation that perturbs the conformational equilibrium, illustrating the assumption that the transition state energy is related to the overall energetic difference of the reaction by a linear factor *ϕ*. (c) Illustration of the 2D manifold defined by the conformational balance equation when varying ΔΔ*G*_*flip*_ or *k*_+_⁄*k*_−_, with *ϕ* fixed to 0.5. (d) Cartoon showing a double-mutant cycle where two mutations push the transporter variants’ conformational balance in the same direction. (e) Double-mutant cycle plot with the functional values from (d), demonstrating how the conformational landscape leads to negative epistasis. The functional score of the WT protein on the left (gray), the two singles in the middle (light blue) and the double mutant in the right (dark blue). (f-g) Similar cartoon (f) and plot (g) for mutations pushing the conformational balance in opposite directions, leading to reciprocal sign epistasis. (h) The same set of mutations as in (f) shown on an altered landscape representing transport of a different substrate with increased baseline *k*_+_⁄*k*_−_. (i) The mutational cycle in specificity space matches the unusual shape of the M230A+E134N double-mutant cycle (Fig. 5c).

This describes a family of bell-shaped curves for the relationship between the ΔΔ*G*_*flip*_ of a variant and its *V*_*max*_ with two free parameters: the transition state *ϕ*-value, which distorts the curve but does not change its overall shape or maximum value (see Supplementary Fig. 5) and the baseline ratio of forward-to-reverse transporter flip rates 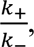 which shifts both the optimal ΔΔ*G*_*flip*_ and the *V*_*max*_ at that optimum (Fig. 7c). As it models the effect of a mutation, we report this equation relative to the WT protein’s maximal rate *V*_*max*,0_. The non-monotonic nature of these curves means that double-mutant cycles of mutants that affect only ΔΔ*G*_*flip*_ can exhibit both negative and positive epistasis, including reciprocal sign epistasis (Fig. 7d-g). Reciprocal sign epistasis is not possible from a monotonic global epistasis function but can be a direct consequence of mutations affecting the Δ*G*_*flip*_ of a transporter. As the M230A mutation has a negative epistatic relationship with many mutations to the outer vestibule—which we hypothesize destabilize the inward-open conformations and bias the transporter toward the outward-open conformation—and a positive epistatic relationship with TM6b mutations—which have been observed to bias the transporter toward the inward-open conformation^40^—this model would predict that M230A should itself push the protein toward the outward-open conformation. This has been observed via cysteine accessibility assay^40^, lending some credence to this model.

This simple family of kinetic curves also provides a plausible mechanism for how the same mutations might affect epistasis and specificity. For a mutation that affects Δ*G*_*flip*_ but not substrate binding, preferential substrate binding to one state in the WT context will affect the baseline *k*_+_⁄*k*_−_ by altering *k*_+_ while *k*_−_ remains the same. It thus will shift the optimal ΔΔ*G*_*flip*_, the overall position of the curve, and the magnitudes or even directions of mutation effects on V_max_ (Fig. 7h-i). If mutations such as E134N indeed bias the protein toward an inward-open state, the fact that they also bias M230A away from Mg^2+^ and toward Mn^2+^ suggests that Mg^2+^ may preferentially bind to the inward-open conformation relative to Mn^2+^. This would explain the overall trend in our results and broadly help explain why M230A has such a uniquely large effect on Mg^2+^ transport for DraNramp but not all canonical Nramps^1,17^. That is, the model suggests that in addition to improving Mg^2+^ binding, M230A brings DraNramp’s baseline conformational equilibrium from the optimal position for Mn^2+^ import to the optimal position for Mg^2+^ import.

The actual biochemistry of DraNramp is likely far more complicated than the toy model described above, but this model provides a first step for understanding why particular mutations could have specificity effects that are highly context-dependent without any obvious mechanism of altering binding directly. Some trends hint at the more complex reality: for instance, the epistatic relationship between H232Y and L381N/C382S seems too specific to relate to the global inward-outward equilibrium and may instead involve a more nuanced conformational change localized to TM10. Nonetheless, our data support a broad model in which metal transport specificity and activity is intimately linked to the inward-outward cycle of transporter function.

## Discussion

Despite advances in determining^46^ and predicting^47^ protein sequence-structure relationships, we still largely lack a predictive biophysical understanding of protein sequence-function relationships. Despite some advances in predicting the activity of very simple biochemical functions such as folding^48,49^ and binding^50,51^, the sequence rules underlying complex phenotypes such as selective transport across membranes that require conformational dynamics remain particularly opaque. In this work, we systematically deconstructed the determinants of metal import and specificity in the model bacterial importer DraNramp by assessing the activity levels of over 30,000 variants in a set of new high-throughput assays for the import of Mn^2+^ or Mg^2+^ to identify principles underlying selective membrane protein transport in this system.

To enrich for informative mutants, we developed a targeted variant library incorporating information from known structures and homologous sequences that allowed us to assess the activity levels both of a broad panel of single mutations and many different combinatorial mutations. We find that DraNramp’s mutational landscape features widespread specific epistasis at non-contacting positions that tend to cluster at positions in the inner and outer vestibules of the transporter. By profiling many mutations on different backgrounds, we identified mutations at some “core” positions—54, 85, 230, 232, and 381—that perturb specificity in favor of Mg^2+^ in all backgrounds and can break WT DraNramp’s efficient exclusion of Mg^2+^, while others are “modulators” that can fine-tune specificity on a background that already transports both ions. These modulator positions overlap those in epistatic hotspots, and we developed a theoretical framework for understanding how mutations that affect the balance between outward- and inward-open conformations could be hotspots for both epistasis and specificity modulation.

The distinction between core and modulatory specificity residues may be a useful framework for understanding specificity evolution in other transporters and selective proteins more broadly. Mutations distal to the binding site are well-known to affect transporter specificity^21,52-54^ but are rarely tested systematically in high throughput. Existing studies profiling the effect on substrate specificity of many transporter variants test only native substrates of the wildtype protein on a single sequence background^22,53,55^. Our study has the advantage of effectively analyzing both adaptation to a non-native substrate and shifting the balance between two existing substrates by profiling mutants of both the Mn^2+^-selective WT background and on the more promiscuous M230A background. The structurally and biochemically diverse profile of variants affecting specificity on the M230A background are more reminiscent of what was recently observed for the multidrug exporter NorA^53^. The contrast with the very limited set of variants that allow for Mg^2+^ import on the WT background highlights how much narrower the paths to new function are than those to modulate existing function, although it is remarkable that multiple single mutations take DraNramp from perfectly selective against Mg^2+^ to robustly able to import enough Mg^2+^ to substitute for *E. coli*’s existing Mg^2+^ importers.

We find that most mutation effects on the native activity of DraNramp, Mn^2+^ import, can be explained by a simple site-independent global epistasis model, but that the effects of mutations allowing for Mg^2+^ import are generally not additive. This contrasts with the yeast pheromone exporter Ste6, for which a wide range of mutations across its binding cavity were found to have small but additive effects on improving transport of a non-native peptide substrate^56^. This likely reflects the difference in substrates: the peptide substrate has a very large interaction surface with Ste6 that is the sum of many contacts, while DraNramp’s interaction with its divalent cation substrates depend on few precisely positioned residues. The additivity of mutation effects in Ste6 also suggests that perhaps the mutations affecting conformational equilibrium in its ABC transporter fold are more segregated from those altering substrate specificity, while in DraNramp these mutations significantly overlap. The distinction between DraNramp and Ste6 highlights how the biochemical natures of different systems shape the structure of their adaptive fitness landscapes.

How distal mutations modulate transport specificity also remains quite opaque despite structural and functional studies. One explanation for soluble proteins is that these mutations promote a new conformation that allows for the new activity^57,58^. In dynamic proteins like transporters that must engage multiple states, a shift in the balance between existing conformations has also been proposed as a mechanism by which distal mutations can alter substrate specificity^22,54^. In our study, the insights we gain from linking specificity modulation to epistasis allow us to build a simple biochemical model of how mutations alter the conformational equilibrium. Although likely overly simple, this model makes several directly testable predictions about which mutations and substrates alter conformational equilibrium between the two states and how this will influence the overall transport rates. These predictions can be expanded in future studies.

Our study also broadly proposes a route to synthesizing two diverging schools of thoughts that have emerged regarding epistasis in protein evolution. Descriptive studies rooted in evolutionary biology have often highlighted the inherent complexity of protein mutational landscapes, replete not only with pairwise epistasis^59^ but also with large numbers of higher-order epistatic terms^24,26,60,61^. Based on this picture, a protein’s fitness landscape might seem too complex to ever possibly predict or fully understand. Alternately, other studies have suggested that protein mutational landscapes can sometimes be far simpler and easier to predict based on simple biophysical modeling^30,38,39,62^. However, a limitation of these studies is that they often use small, model proteins and fit to simple phenotypes such as folding and binding.

Selective transport represents a more complex phenotype with many different biochemical parameters. Thus it is unsurprising that we find more complex epistasis patterns in the fitness landscape of DraNramp for Mn^2+^ import than has been observed in systematic studies of protein folding, where pairwise epistasis that cannot be explained by a monotonic global epistasis function primarily occurs at structural contacts^30^. However, we identify considerable structure in the residuals beyond the simple baseline assumptions. Mutations with different classes of deviations from the null expectation tend to structurally cluster, and, strikingly, the residuals from two independent analyses of epistasis and specificity strongly anticorrelate, suggesting that they are largely shaped by a core mechanism. This opens the door to identifying predictable structure in the complexity of DraNramp’s mutational landscape. Our conformational balance model explains some of the variability in this landscape, but fully explaining the complex structure is still limited by the sparse scale and type of data we collected.

Generating the data to build the next generation of models linking transporter biochemistry to evolution will require new innovations. First, with the large number of mutations in our evolution-guided library, we could not measure the effect of each mutation on enough backgrounds to fit more complex models. More densely sampling combinations of mutations would measure the effect of every mutation and mutation pairs on many backgrounds, allowing for a more systematic and quantitative analysis of epistasis. However, even with very large mutational libraries, it may be challenging to distinguish between competing biophysical models. To address this, it would be valuable to make additional orthogonal measurements for the same library such as expression levels, dose-response curves, or even reading out the conformational populations. Further, assays conducted in cells typically come with various artifacts that complicate our understanding of fundamental protein biophysics. Recent developments in high-throughput biophysics done *in vitro*^63,64^ are currently limited to soluble domains but could in the future be adapted for membrane protein systems to do high-throughput transport biochemistry.

In summary, we have used a large, targeted library of single and combinatorial variants alongside new high-throughput assays to profile the determinants of import for both a native and non-native substrate to the model bacterial metal importer DraNramp. We identified a set of core specificity residues that have the largest effects on enabling DraNramp to transport Mg^2+^ and an additional set of modulator mutations that overlap considerably with hotspots of epistasis in the fitness landscape for DraNramp’s native substrate of Mn^2+^. These results allow us to propose new models for how transporter thermodynamics can shape their fitness landscapes and broadly provide a framework for future work developing a quantitative understanding of sequence-function relationships for dynamic proteins.

## Methods

### Binding-site library design

The binding-site library consists of mutations at every position within 12 Å of the central Mn^2+^ ion in any of three high-resolution metal-bound DraNramp structures (PDB IDs: 8E6N, 8E60, or 8E6I). For each position, we mutated the relevant codon to the most abundant *E. coli* codon for each amino acid, as well as to the most abundant non-WT synonymous codon (if possible) and deleted the codon entirely. These sequences were ordered as one of two oligo pools from Twist Biosciences (Supplementary Table 4). Each subpool was individually assembled into pET28a by Gibson assembly, adding an 18-bp barcode of sequence NNNNSWNNNNSWNNNNSW^65^ to the 3’ multiple cloning site. We transformed these ligation products into electrocompetent DH10B *E. coli* (GoldBio) and the requisite number of colonies were harvested from agar plates to bottleneck each subpool to ∼40X the subpool diversity to balance obtaining nearly every variant with attaining a reasonable barcode diversity. Subpool plasmid DNA was isolated using a ZymoPURE II Plasmid Midiprep Kit and pooled in appropriate molar ratios to balance subpool diversity.

### Building generative sequence models

We used our recently described MSA of Nramps and Nramp-related proteins^66^ as an input for all sequence models. This MSA contains 17,763 sequences from across the tree of life filtered to 90% sequence identity, all of which are more similar evolutionarily to DraNramp than to the sodium-neurotransmitter symporter family and thus likely metal transporters (Extended Data Fig. 1a). Multiple metal ion substrate specificity profiles are represented, although the specificity profile of most Nramp-related proteins in this alignment is unknown. We ran EVcouplings v0.2.1 using the standard pipeline with inference via pseudolikelihood inference, a minimum sequence coverage of 50%, a minimum column coverage of 70%, and *θ* = 0.8. This pipeline generates both an “epistatic” EVcouplings model and a site-independent position-specific scoring matrix (PSSM). We verified that the top couplings (>90% probability) produced from this model closely match structural contacts in either of DraNramp’s inward-open or outward-open conformations (Extended Data Fig. 1b). Sequence logos in Fig. 1b and Extended Data Fig. 1e were generated using our recently described tree^66^ and visualized with logomaker v0.8 in Python^67^, with clades selected manually by partitioning the tree and discarding very small clades with fewer than 50 sequences.

### Gibbs sampling of protein sequences from arbitrary fitness models

To sample from a given probability distribution over sequence space *p*(*x*_*i*_), we use Gibbs sampling by first iterating through each sequence position in a random order and, at each position, selecting a new amino acid from the distribution *p*(*x*_*i*_|*x*_*i*≠*j*_). For energy-based models such as EVmutation, *p*(*x*_*i*_) is related to the underlying computational “energy” of a sequence *U*(*x*) via the Boltzmann distribution, i.e. 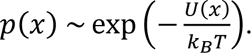 For all sampling, we set the computational temperature *T* by screening a range of temperatures and selecting one that generated sequences with a similar average energy to WT DraNramp. To sample sequences close in Hamming distance to WT DraNramp, we also added a distance restraint to the energy function *U*(*x*), such that:

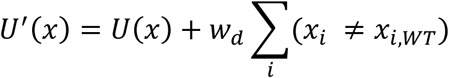

where *w*_*d*_ is an arbitrary parameter that controls the strength of the distance restraint (Extended Data Fig. 1c). The linear form of this restraint approximately cancels out the increasing entropy of sampling sequences more distant from a reference.

### Latent exploration sampling

To sample a set of sequences predicted to be enriched for specificity-switching variants, we first constructed a lower-dimensional sequence space representation of the alignment using principal component analysis (PCA) in scikit-learn v1.0.8 ^68^. This decomposes the one-hot encoded alignment X into three matrices *U*, Σ, and *V* (Fig.1c), in which the columns of *U* represent embeddings of each sequence in the alignment in the PCA space and the columns of *V* represent orthogonal vectors in mutation space^31^. When partitioning the alignment into discrete clades, mutations with the largest weights in the principal components 𝑣 are generally conserved within clades but not between clades. These mutations are enriched around the binding site although some mutations are distributed throughout the protein core. Notably, this approach is similar to statistical coupling analysis (SCA)^69^, which uses a conservation-weighted version of PCA to decompose the alignment. However, due to how it implements the decomposition SCA can only suggest positions and not individual amino acid changes at those positions. Each possible variant of DraNramp will alter its embedding *u*_*i*_, and those that include the mutations with the largest weights in 𝑣^(1)^ and 𝑣^(2)^ will alter it most. We hypothesized that if this sequence space representation encodes any functional information, these mutations should be more likely to alter, rather than abolish, function. We therefore included a linear bias term into the energy-based sampling framework introduced above to decrease the computational energy of a sequence based on its distance from the WT in the embedding space as an “exploration potential.” Mathematically, this modifies the underlying energy term (or logit) of any generative sequence model as follows:

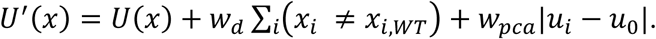

Here, the *w*_*d*_ ∑_*i*_(*x*_*i*_ ≠ *x*_*i*,*WT*_) term adjusts the model’s distance restraint to sample sequences with few absolute mutations relative to a reference sequence (*x*_*WT*_; a “pull” toward WT in sequence space) while the *w*_*pca*_|*z*_*i*_ − *z*_0_| term rewards sequences for changing their latent embedding (a “push” away from WT in functional space). For the Nramp evolution-guided library, we spanned a range of values of *w*_*d*_ and *w*_*p*_ and ultimately selected *w*_*d*_ = 8.25 and *w*_*p*_ = 40 to match our desired distance distribution peaking at 2-4 mutations and maximizing the sampling of known positive controls mutation M230 without increasing the frequency of the D56G mutation beyond one-third of the library (Extended Data Fig. 1d).

### Evolution-guided library construction

To convert samples from the generative latent space exploration model (the “design model”) described above into physical DNA (the “evolution-guided library”), we used an approach inspired by variational synthesis^32^. The concept is to approximate sampling from the probability distribution of a generative sequence model under the constraints of a stochastic synthesis protocol (the “stochastic synthesis protocol”). This was necessary as the length of DraNramp made ordering exact combinatorial variants (for example, on a single oligo pool) impossible. We chose as our stochastic synthesis protocol a “spread-out low-diversity library” from Twist Biosciences, which allowed us to define arbitrary codon frequencies at up to 60 positions across a region of approximately 1000 bp. Notably, while variational synthesis requires the generative synthesis protocol to have support over all possible sequences, spread-out low diversity libraries do not meet this criterion as most positions are fixed. The optimal design was thus given via a two-step process: first, we selected the subset of 59 positions most mutated by the design model; then, at these positions, we optimized the frequencies of the library design to match the design model. The library synthesis protocol in Supplementary Table 5 was used to synthesize the library based on a codon-optimized Nramp sequence.

We assembled this linear library fragment into a pET28a backbone using Gibson assembly and transformed via electroporation into DH10B *E. coli*. Cells were recovered in lysogeny broth (LB) for 1 hour, then different dilutions were added to six separate flasks of 25 mL LB supplemented with 50 µg/mL kanamycin and 0.35% SeaPrep soft agarose. This soft agarose mixture was set at 4°C for 2 hours before growing at 37°C for 12 hours to allow for the formation of individual agarose-embedded colonies, which promotes more even growth of transformants after recovery. We blended each soft agarose gel by shaking for 1 hour at 37°C, then diluted 1:50 into 50 mL LB to grow overnight at 37°C in liquid culture before isolating plasmid DNA via midiprep (Zymogen).

To assess the number of unique barcodes in each library, we performed small-scale Illumina sequencing on the amplified barcodes of each sample via Amplicon-EZ (Azenta), obtaining between 120,000 and 220,000 reads per sample. We discarded barcodes observed only once as likely sequencing errors and then counted the total number of barcodes in each library. Based on this analysis, we pooled the four of the six input libraries into a final library with an estimated diversity of >45,000 unique barcodes.

### Barcode-variant mapping

To map individual barcodes from the library to variants, we digested the DraNramp gene from the plasmid library with EcoRI-HF and HindIII-HF, gel-purified it, and sequenced the amplicons via Nanopore (Plasmidsaurus), yielding 6,878,645 and 8,539,764 raw reads for the binding-site and evolution-guided libraries respectively, which we aligned to the expected wildtype library sequence with Minimap v2.1^70^. We sorted the reads by barcode using the python package pysam, then generated a consensus sequence for all reads sharing the same barcode using medaka (Oxford Nanopore).

### Cloning individual variants

Individual variants were cloned out of the library via a two-step process. First, the variant of interest was amplified from 10 ng of pooled library with a reverse primer matching its unique barcode and a forward primer at the 5’-end of the DraNramp gene. These PCR products were gel-purified and used as templates for a second PCR reaction with longer primers allowing for Gibson assembly into a pET28a backbone digested by EcoRI and HindIII. All primer sequences are included in Supplementary Table 6. These assembly products were transformed into DH5*α* cells and plated on LB supplemented with 50 mg/mL kanamycin. Plasmids were isolated from single colonies, and sequences were verified with whole-plasmid sequencing (Plasmidsaurus).

### Creation of recombinant-deficient E. coli strains and competent cells

We knocked out the *E. coli* recA gene in both OverExpress C41(DE3) and the BW25113 Δ*mgtA* Δ*corA* Δ*yhiD* DE3 strain^44^ (gift of Osamu Nureki and Motoyoki Hattori) using P1 phage transduction from the recA knockout in the Keio collection^71^, to create the C41Δ*recA* and the MgKO strains, respectively. The use of C41Δ*recA* rather than other *E. coli* strains pre-optimized for screening was essential due to the ability of C41Δ*recA* cells to express much higher levels of the mildly toxic DraNramp protein. We then looped out the kanamycin resistance cassette by transforming the pCP20 plasmid (gift of Daniel Kahne) into these cells and growing overnight at 42°C to promote plasmid loss. We selected colonies that grew on LB alone but not LB supplemented with 50 µg/mL kanamycin and confirmed each knockout strain via colony PCR and Nanopore sequencing (Plasmidsaurus).

We prepared chemically and electrocompetent cells using standard protocols. Briefly, chemically-competent C41Δ*recA* and pSB1-C41Δ*recA* were prepared from single colonies expanded in 200 mL LB (with 25 µg/mL chloramphenicol for pSB1-C41Δ*recA*) to OD_600_ ∼ 0.5. Cells were pelleted and resuspended in 2-3 mL of Solution I (70 mM CaCl_2_, 50 mM MgCl_2_), incubated on ice for an hour, and then an equal volume of Solution II (70 mM CaCl_2_, 50 mM MgCl_2_, 50% glycerol) was added. Electrocompetent pSB1-C41*ΔrecA* cells were prepared expanding a single colony in 1L LB with 25 µg/mL chloramphenicol to OD_600_ ∼ 0.5. Cells were washed in 1, 0.5, and then 0.25 L of ice-cold sterile 10% glycerol before resuspending in 10% glycerol at OD_600_ ∼ 100 (∼1-2 mL). Electrocompetent MgKO-Δ*recA* were prepared similarly, but the initial growth was done in LB supplemented with 100 mM MgSO_4_. Electrocompetent cells were flash frozen and stored at -80°C.

### Cloning pSB1 plasmids

The Mn^2+^ reporter plasmid was constructed in three pieces. The noncoding region between the genes *yopD* and *mntP* containing the *mntP* promoter and the *yybP*-*ykoY* riboswitch was amplified from Nissle 1917 *E. coli* genomic DNA. The gene encoding mEmerald was amplified from the pEJM25 plasmid (gift of Dr. Elizabeth May). For pSB1, both were inserted via three-piece Gibson assembly into the backbone of a pACYC-1-Duet plasmid digested with EcoNI and KpnI. For pSB1-EV, only the *mntP* 5’ region but not mEmerald was inserted via two-piece Gibson assembly. Plasmid sequences were confirmed via whole-plasmid sequencing (Plasmidsaurus) and can be found in Supplementary Table 6. We transformed pSB1 and pSB1-EV into C41Δ*recA* cells to created pSB1-C41Δ*recA* and pSB1-EV-C41Δ*recA*, respectively.

### Mn^2+^ import assay on single variants

We chemically transformed pET28a-DraNramp variants into pSB1-C41Δ*recA* cells, and picked single colonies from LB-agar plates supplemented with 50 µg/mL kanamycin and 25 µg/mL chloramphenicol into 5 mL overnight cultures of rich defined assay medium (RDAM; 33.7 mM Na_2_HPO_4_, 22 mM KH_2_PO_4_, 8.55 mM NaCl, 9.35 mM NH_4_Cl, 1 mM MgSO_4_, 0.3 mM CaCl_2_, 134 µM EDTA, 31 µM FeCl_3_, 6.2 µM ZnCl_2_, 0.76 µM CuCl_2_, 0.42 µM CoCl_2_, 1.62 µM H_3_BO_3_, 1 µg/mL thiamin supplemented with 0.4% glucose, 0.25% casamino acids, 50 µg/mL kanamycin and 25 µg/mL chloramphenicol). RDAM is based on M9 medium with a defined elemental composition to exclude Mn^2+^, which is present in low amounts in most standard media and tap water. Saturated overnight cultures were diluted 1:50 into 5 mL RDAM. For the plate reader assay shown in Extended Data Fig. 2a, 190 µL of the diluted culture was added to a black clear-bottom 96-well plate (Greiner) and grown on a Biotek Neo2 plate reader at 37°C with shaking. After cultures reached an OD_600_ = 0.4, they were induced with 250 µM IPTG, and then after 30 minutes we added varying amounts of MnCl_2_. mEmerald fluorescence was measured with an excitation wavelength of 479 nm and emission wavelength of 520 nm. Growth curves were analyzed using a custom script in Python and normalized against the fluorescence values for pSB1-EV-carrying cells, approximately correcting for the background fluorescence increase due to increasing cell density.

For the flow cytometry-based assay (as in Fig. 2b-c), cultures were initially grown in 5 mL RDAM through induction, then 190 µL of the culture was transferred to a 96-well plate just before MnCl_2_ addition and grown at 37°C with shaking for four hours. Cultures, at an OD_600_ ∼ 2.0, were diluted 40-fold to an OD_600_ ∼ 0.1 for flow cytometry. Forward scatter, side scatter, and green fluorescence (ex. 480 nm / em. 520 nm) were recorded for 10,000 events for each well on a FACS Aria Symphony equipped with a high-throughput sampler (HTS). Raw .fcs files were analyzed in python using the flowkit package. In many cases, cells have a multimodal distribution when plotted in SSC-A/FITC-A space (Extended Data Fig. 2b). We hypothesize that some cells silence transporter expression, as the smaller population approximately matches that of the inactive D56A control at the given Mn^2+^ concentration. We modeled the distributions as a two-component Gaussian mixture model where one component is regularized to match an “inactive” population fit to the negative control population. For each Mn^2+^ concentration, we first fit a single two-dimensional Gaussian with mean *μ*_*inactive*_ and covariance Σ_*inactive*_ to the log(FITC-A) and SSC-A values for the negative control sample. Then we fit a two-component Gaussian mixture model to all other samples where the first (“inactive”) component is heavily regularized to match the “inactive” distribution and to have a small total fraction. We do so using a standard expectation-maximization (E-M) algorithm, with a standard E-step and an M step in which we optimize the regularized log-likelihood:

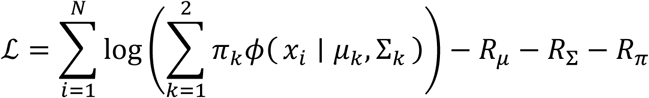

Where the three regularization terms are given by:

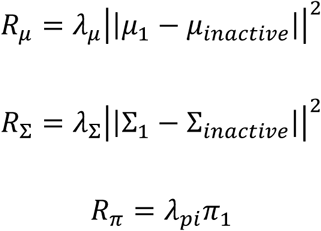

The regularization weights were tuned empirically, with final values of *λ*_*μ*_ = 0.3, *λ*_Σ_ = 0.3 and *λ*_π_ = 0.1. After convergence, we then took the FITC-A mean of the second Gaussian for each sample as the mean log fluorescence score for the active population of cells. Only single Gaussians were fit for all negative control populations, including both cells with an empty vector and those overexpressing the inactive variant D56A.

To generate normalized dose-response curves, the mean log fluorescence score of D56A for the relevant Mn^2+^ concentration was subtracted from each data point. We chose D56A rather than empty vector as D56A had consistently lower fluorescence scores at high [Mn^2+^], likely due to the fitness cost on the cell imposed by protein overexpression. Because all other samples were overexpressed protein and thus likely had a similar fitness cost, this was the most comparable negative control to use as a baseline. We fit a Michaelis-Menten curve to each of these dose-responses using the curve_fit function in sklearn v1.0.2 with an initial guess (*p*_0_) of 0.1 for the V_max_ and 25 for the K_M_.

### Mn^2+^ import screen

The day before the screen, 10 ng of the relevant pET28a-DraNramp plasmid library was electroporated into each of two 100 µL aliquots of electrocompetent pSB1-C41Δ*recA* cells. Clonal stocks of WT DraNramp, N59D, and D56A were also electroporated in parallel. After 1 hour of recovery at 32°C in SOC medium, library transformations were added to 225 mL 0.35% SeaPrep soft agarose in RDAM, while control transformations were added to 15 mL of the same mixture. These mixtures were cooled at 4°C for two hours, then grown for 12 hours at 37°C. In parallel, we estimated the transformation efficiency by plating serial dilutions of cells in triplicate after recovery. We obtained more than 10 million transformed colonies per replicate, surpassing 100X coverage for each library.

On the day of the screen, the soft agar colony suspensions were blended by shaking at 37°C for 30 minutes and then a volume approximating a final 1:50 dilution of a culture at OD_600_ = 2.0 was added to 10 mL RDAM. Cells were induced at OD_600_ = 0.4 and protein expression was induced with 250 µM IPTG. After 30 minutes, 100 µM MnCl_2_ was added and cultures were grown with shaking at 37°C for 4 hours. One mL of each culture was removed and washed twice with 50 mL 1X M9 medium (33.7 mM Na_2_HPO_4_, 22 mM KH_2_PO_4_, 8.55 mM NaCl, 9.35 mM NH_4_Cl) to remove residual Mn^2+^ and agar. Cells were then passed through a sterile 40 µm cell strainer and diluted up to OD_600_ = 0.1 (∼10^8^ cells/mL) in 1X M9 medium.

Cells were sorted into four bins on a FACS Aria II. On each screen day the gates were adjusted to match the approximate gate frequencies of the controls for the first replicate. Cells were first gated for size on FSC-A / SSC-A and then on SSC-W / SSC-H to select against doublets (Extended Data Fig. 3a). Final gates were then set in two dimensions on the FITC-A / SSC-A plot, as FITC-A correlates positively with SSC-A and thus setting gates that follow this correlation trend allows for better separation between controls, and thus likely between variants in the library (Extended Data Fig. 3b). Greater than 15 million cells were collected across the four gates for each replicate of the evolution-guided library screen, and greater than 5 million cells per replicate for the smaller binding-site library. For the evolution-guided library, each of four replicates was performed on a separate day, while the three binding-site library replicates were done in parallel. As we observed significant bias in growth between different control variants when recovering cells overnight after sorting, we elected to directly collect DNA from sorted cells. To minimize plasmid loss on miniprep columns optimized for larger volumes of cells, we determined that adding additional pSB1-C41Δ*recA* cells containing the pSB1 reporter but no pET28a plasmid improved yield of library plasmid from small numbers of cells. To improve cell and plasmid harvesting, we added independently grown pSB1-C41Δ*recA* cells corresponding to ∼100X the number of cells sorted into the largest gate before immediately pelleting sorted cells and extracting DNA via standard miniprep (Omega).

### Mg^2+^ import assay on individual variants

The relevant expression plasmid was electroporated into MgKO cells and cells recovered in 100 mM MgSO_4_ for two hours at 37°C before plating on LB supplemented with 50 µg/mL kanamycin (LB-kan) and 100 mM MgSO_4_. Single colonies were grown overnight in LB-kan with 100 mM MgSO_4_, then normalized to OD_600_ = 2. Cells were diluted 1:100 into LB-kan and pipetted into a clear flat bottom 96-well plate (190 µL per well). Cell growth was measured at different concentrations of Mg^2+^ and IPTG by adding 5 µL of 40x MgCl_2_ and 40x IPTG solutions to each well. Cultures were shaken at 37°C for 12 hours and cell density (OD_600_) measured every two minutes in a BioTek Epoch 2 microplate spectrophotometer.

To analyze growth curves, we used the Python package Curveball which fits various growth models to growth curve data^72^. Curveball selects the best-fitting model from a nested family of growth models, ranging from the three-parameter logistic model to the six-parameter Baranyi-Roberts model^73^, using the Bayesian Information Criterion (BIC). In some samples, we observed biphasic growth curves, characterized by an initial growth phase followed by a variable-length lag phase, after which a second distinct growth phase emerged. This biphasic behavior resulted in poor model fits for some growth curves. To address this, we restricted the dataset to the first 8 hours of growth and fit the models only to the initial growth phase to extract parameters. To further reduce spurious fits, we constrained the initial population *y*_0_ between 0 and 0.05, the saturation population size *K* to between 0.4 and 10, the exponential rate *r* to be positive, the logistic curvature parameter *v* to be greater than 1, and the Baranyi-Roberts lag parameters *q*_0_ and *v* to each be greater than 0.01. For subsequent analysis, we report only the growth rate (*r*) for the first growth phase.

### Mg^2+^ import screen

Each variant library was independently transformed into MgKO cells via electroporation, with 10 ng DNA added to each 100 µL aliquot of competent cells. Cells were grown overnight at 37°C in LB-kan with 100 mM MgSO_4_ supplemented with 0.35% SeaPrep agarose. Embedded colonies were blended via shaking at 37°C until OD_600_ ∼ 2.0, then cells were diluted 1:100 into LB-kan in glass round bottom tubes to a final volume of 10 mL, and then supplemented to a final concentration of 1, 5, or 100 mM Mg^2+^. DNA from a “pre-selection” population of cells was collected at this stage. For all three Mg^2+^ concentrations, protein expression was induced with 20 µM IPTG. An additional condition included no induced expression (i.e., no IPTG) and 1 mM Mg^2+^. Cultures were shaken at 37°C for 12 hours after which the DNA was isolated from each sample via standard miniprep (Omega).

### Illumina sequencing

To generate barcoded amplicons for Illumina sequencing, we performed two rounds of PCR on the raw sorted or selected plasmid libraries with a TruSeq-based primer design strategies adapted from Lue et al. (2023)^74^. In the first round of PCRs, we added inline offsets and TruSeq adapter regions. In our first sequencing runs (replicates 1 and 2 of the Mn^2+^ screen for the evolution-guided library), we also included unique molecular identifiers (UMIs) in these first-round primers (random 8-bp stretches between the plasmid binding region and TruSeq adapter) and ran this first-round reaction for only 3 cycles before digesting the original template with Dpn1 and isolating DNA via PCR purification. Although the UMI allows for downstream PCR deduplication by counting unique UMIs per variant barcode, this deduplication did not meaningfully change our results. For later PCR reactions we eliminated the UMIs and opted for 20 cycles on the first-round PCR followed by extraction of the correct-sized amplicon from a 2% agarose gel, which led to far cleaner and more consistent final amplicons. In the second round of PCR, we added appropriate i5 and i7 sample indices for each sample (see Supplementary Table 6 for all primers used for each sample) along with Illumina adapters. All PCRs were done with Q5 polymerase at an annealing temperature of 65°C with a 20-second extension time. The final samples were again extracted from a 2% agarose gel, dsDNA concentration was quantified via Qubit, then samples were appropriately pooled and sequenced.

Sample results are ultimately taken from three separate Illumina sequencing runs done at Harvard’s Bauer Core: a 60x40 cycle NextSeq 1000 run done at Harvard’s Bauer Core and 2 separate lanes on a 100x50 cycle NovaSeqX. Forward and reverse reads were then merged using BBmerge version 38.06^75^ and the counts of each barcode were calculated from each sample with a custom python script.

### Mn^2+^ assay data analysis

For the Mn^2+^ screen data, we first estimated the overall frequency of each variant in each bin by normalizing the total observed counts (*c*_*obs*_) to the overall counts. We assume that the counts *c*_*i*_ for each variant *i* were sampled following a Poisson distribution, *p*(*c*_*i*_|*λ*_*i*_) = *Poisson*(*λ*_*i*_), where *λ*_*i*_ = *f*_*i*_*N*_*reads*_. Here, *f*_*i*_ represents the true underlying frequency of variant *i* in the sample and *N*_*reads*_ were the total number of reads for that sample. To estimate the posterior *p*(*λ*_*i*_|*c*_*i*_), we assumed the prior distribution:

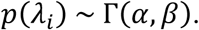

Based on which the posterior distribution will be:

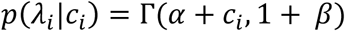

We set *α* = 1 and *β* = 0.001 as a generally uninformative prior that nonetheless estimates appropriately larger errors for samples with few read counts. To estimate *f*_*i*_, we first took the expected value of this distribution:

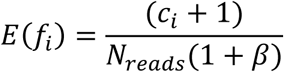

Given the frequencies in each bin for each variant, we can calculate a histogram (Extended Data Fig. 3d) that approximates the original fluorescence distribution by bin. To convert overall frequencies among reads to estimated fraction of sorted cells, we first normalized by the number of cells sorted into each bin. As a summary metric, we calculated the weighted mean bin of these histograms and normalized it such that the WT protein has a score of 1 and a representative inactive frameshift at position 3 has a score of 0. We estimated an error distribution on this by drawing 100 samples from *p*(*λ*_*i*_|*c*_*i*_) and calculating a distribution of possible scores for the variant given the read count error, which we summarize with a standard deviation. For final scores, we merged observations between replicates (*n=*4 for the evolution-guided library and *n*=3 for the binding-site library) using the random effects model from enrich2 to combine the estimated read count error with the replicate error^76^.

### Mg^2+^ assay data analysis

For the Mg^2+^ assay, we estimated the variant frequency *f*_*i*_ in each condition as above, but rather than calculating a weighted mean bin, we calculated a log enrichment score by dividing *f*_*i*_ for each condition by the frequency of that variant in the pre-selection library. Considerable growth variation in the pre-selection overnights in this assay meant that many variants dropped out of individual replicates, and—despite very deep sequencing—we found that variants with low counts in the pre-selection population for a given replicate had very unreliable scores compared to other replicates. Thus, when calculating mean scores and errors across variants, we ignored replicates for each variant with a pre-selection read count below a given threshold: 50 for the evolution-guided library, 100 for the full pool of the binding-site library, and 5 for the repeat of the TM3+TM6 subpools. This lower threshold for the TM3+TM6 subpool was due to a lower overall number of reads for this pool.

### Library score merging

To merge scores between libraries and generate scores for the “combined library,” we performed a linear regression of overlapping scores between libraries under the assumption that mutational scores in each library should be similar but the scaling of exact scores (and thus the slope of the regression line) could be different. We first filtered scores from individual libraries based on their estimated error such as not to bias the regression based on poorly estimated variants; this threshold was *σ* = 0.15 for each library in the Mn^2+^ assay; *σ* = 0.5 for the binding-site library in the Mg^2+^ assay, and *σ* = 1.0 for the evolution-guided library in the Mg^2+^ assay, for which scores are more spread overall. We then fit a linear model to the read counts from both libraries with orthogonal distance regression (ODR) as implemented in scipy v1.14.1 to model error on the scores from both libraries simultaneously, rather than treating one score as an error-free independent variable (Extended Data Fig. 3i, Extended Data Fig. 6e-d). For the Mg^2+^ data, our repeat of the assay on the binding-site library with the TM3+TM6 subpools only necessitated first merging these scores with the rest of the binding-site library scores. For this regression, the best fit was given with *m* = 1.284 and *b* = 0.2. To merge these scores, we rescaled each of the three individual replicate log enrichment scores from the TM3+TM6 binding site pool with these regression parameters to fit the rest of the binding-site library and then re-estimated the mean and error of the replicates passing their respective read-count thresholds using the Enrich2 random effects model^76^. We then used these merged scores to compare against the evolution-guided library to maximize overlap. Between these libraries, we found the best fit was given with *m* = 0.41 and *b* = −0.16. This indicates that the raw scores from the binding-site library are smaller in magnitude than those from the evolution-guided library, which matches our expectation based on the low fraction of Mg^2+^-transporting variants in the evolution-guided library. The paucity of overlapping variants with moderate values makes it challenging to estimate whether the relationship between scores in these libraries is truly linear, and thus direct comparison of absolute magnitudes of variants tested in these separate libraries should be done with caution.

### Determining significant Mg^2+^ transporters

In the Mg^2+^ dataset, the rarity of M230-retaining variants that rescued MgKO growth made us concerned that some detected hits were truly noise. To assess this possibility, we began by analyzing the distribution of Mg^2+^ import scores among barcodes mapping to WT DraNramp in each sequenced pool (the evolution-guided library, the full binding-site library, and the repeat of TM3+TM6 subpool library; Extended Data Fig. 6f-h). We propagated this uncertainty with the estimated error on the Mg^2+^ import score of each M230-retaining variant to calculate an initial p-value representing the likelihood of measuring this score for a variant with WT-like activity levels. We excluded positions containing M230 mutations from this analysis as these mutations do not share the null hypothesis. We then calculated a false discovery rate that accounts for the total number of hypotheses tested with the Bejamini-Hochberg correction as implemented in the python package statsmodels v0.13.5. Variants with corrected p-values under 0.05 were thus deemed significant, resulting in a total of 37 significant variants lacking M230X mutations.

## Acknowledgements

We acknowledge members of the Gaudet and Marks Labs for useful discussions. We would also like to especially thank members of Harvard’s Bauer core for assistance with cell sorting; Brian Shim, James Woods, Simon Shen, and Claire Reardon for help with Illumina sequencing; Sebastian Rowe for help with phage transduction; Jeffrey Chang for general advice on pooled screening; and Emma Garcia for scientific advice, troubleshooting, and support. This work was funded in part by NIH grant R01GM120996 (R.G.), the NSF-Simons Center for Mathematical and Statistical Analysis of Biology at Harvard (award number 1764269) and the Harvard Quantitative Biology Initiative (S.P.B.), NIH/NIGMS T32 grant GM0008313 (S.P.B., PI Venkatesh N. Murthy), and the Chan Zuckerberg Initiative grant 5R01CA260415 (D.S.M.).

## Author Contributions

S.P.B, D.S.M. and R.G. conceptualized the project. S.P.B. designed and cloned sequence libraries.

S.P.B. and C.B.F. developed assays, performed screens, and followed up on individual variants.

S.P.B. analyzed screen data and performed statistical analyses.

S.P.B., C.B.F. and R.G. wrote the manuscript. All authors reviewed and edited the manuscript.

---

**Extended Data Figure 1.**
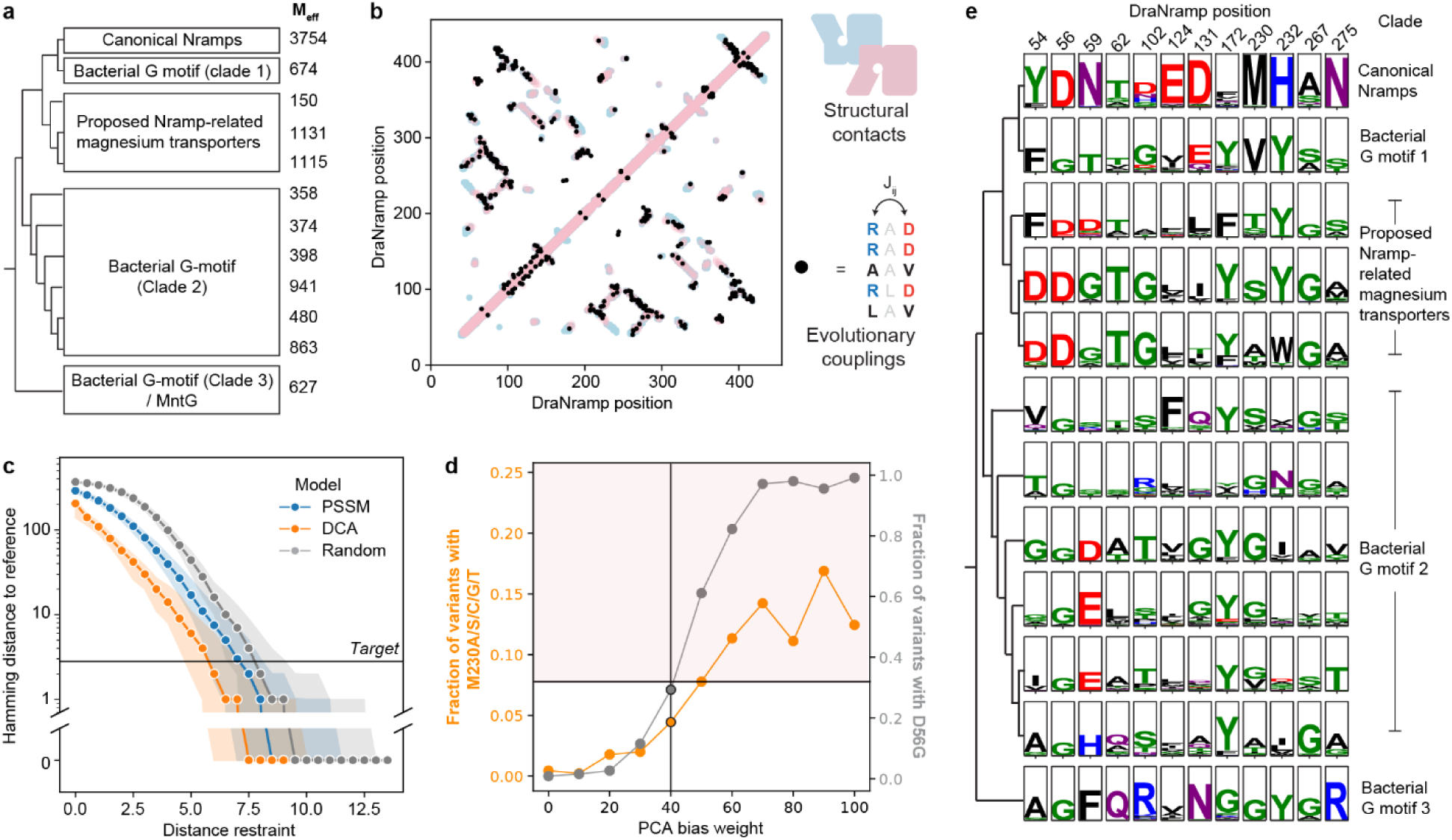
Using evolutionary information to guide sampling of a sequence library. (a) Schematic of the phylogenetic tree of 17,763 aligned sequences of Nramps and Nramp-like transporters, with the effective number of sequences (M_eff_, defined with *θ* = 0.8) per subclade indicated on the right. (b) Overlay of the contact maps for the outward-open (PDB ID: 8E6N, blue) and inward-open (PDB ID: 8E6M, pink) structures of DraNramp with evolutionary couplings (black) predicted based on our 17,763-sequence MSA via the EVcouplings model. Couplings align remarkably well with structural contacts, indicating the alignment is high quality. (c) The distribution of Hamming distances to the WT reference sequence (DraNramp) decays for higher values of the distance restraint when sampling random sequences (gray), from a site-independent model (PSSM; blue) or from an EVcouplings DCA model (orange) with Gibbs sampling. We set the distance restraint for each model to target an average Hamming distance to WT DraNramp of 2.8 (black horizontal line). Ninety-five percent of sampled sequences for a given distance restraint fall within the shaded interval, indicating how for low values of the distance restraint no sequences are ever sampled close to the reference sequence for any of the models. (d) Graph illustrating for two examples of how the PCA bias weight affects how heavily the model samples mutations at differentially conserved positions. For each PCA bias weight, we selected the appropriate distance restraint based on a broad sweep to obtain an average Hamming distance to WT DraNramp of 2.8. The graph shows the fraction of variants from 10,000 samples with positive control mutations (M230A/S/C/G/T; orange, left axis) or with the most heavily weighted mutation (D56G; gray; right axis). Note that the D56G scale is amplified by 4X relative to the M230A/S/C/G/T mutations, as more mutations are sampled with D56G, maxing out at 100% for high PCA bias weights. The black vertical line marks the chosen PCA bias weight for Figure 1c. (e) Sequence motifs from the full alignment for positions with mutations that contribute most to the principal components, as highlighted in Figure 1d. The canonical Nramps are at the top, while different clades (matching those in panel a and Figure 1b) are separated out with a schematized tree on the left. Sequence logos were generated with logomaker in python and letter heights represent information in bits.

**Extended Data Figure 2.**
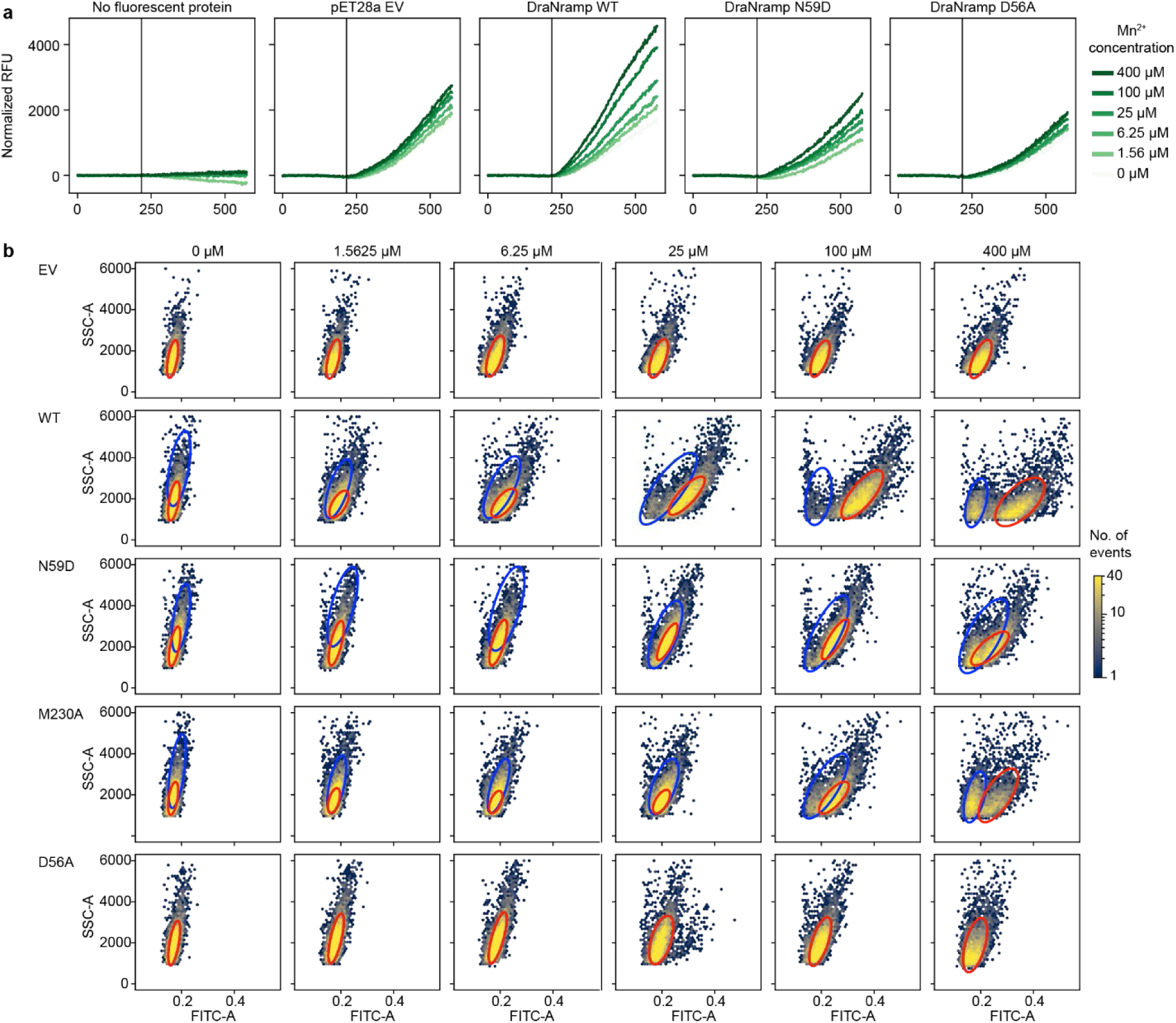
Example data from the Mn^2+^ import assay. (a) Bulk fluorescence over time for pSB1-C41Δ*recA* cells overexpressing different DraNramp constructs, with an “empty” pSB1-EV reporter plasmid lacking the fluorescent protein coding sequence on the far left. Different colored curves represent different MnCl_2_ concentrations. (b) Distributions of green fluorescence (FITC-A) measurements against approximate cell size measured by side scatter (SSC-A) on a flow cytometer for different DraNramp constructs in pSB1-C41Δ*recA* and different MnCl_2_ concentrations. Particularly at high [MnCl_2_], cells are somewhat bimodal. Distributions of core “active” (red ovals) and “inactive” (blue ovals) populations were identified by the structured Gaussian mixture model (see Methods). The FITC-A means of the red (“active”) distributions were used for downstream analyses. The color on each plot represents the density of events within a bin.

**Extended Data Figure 3.**
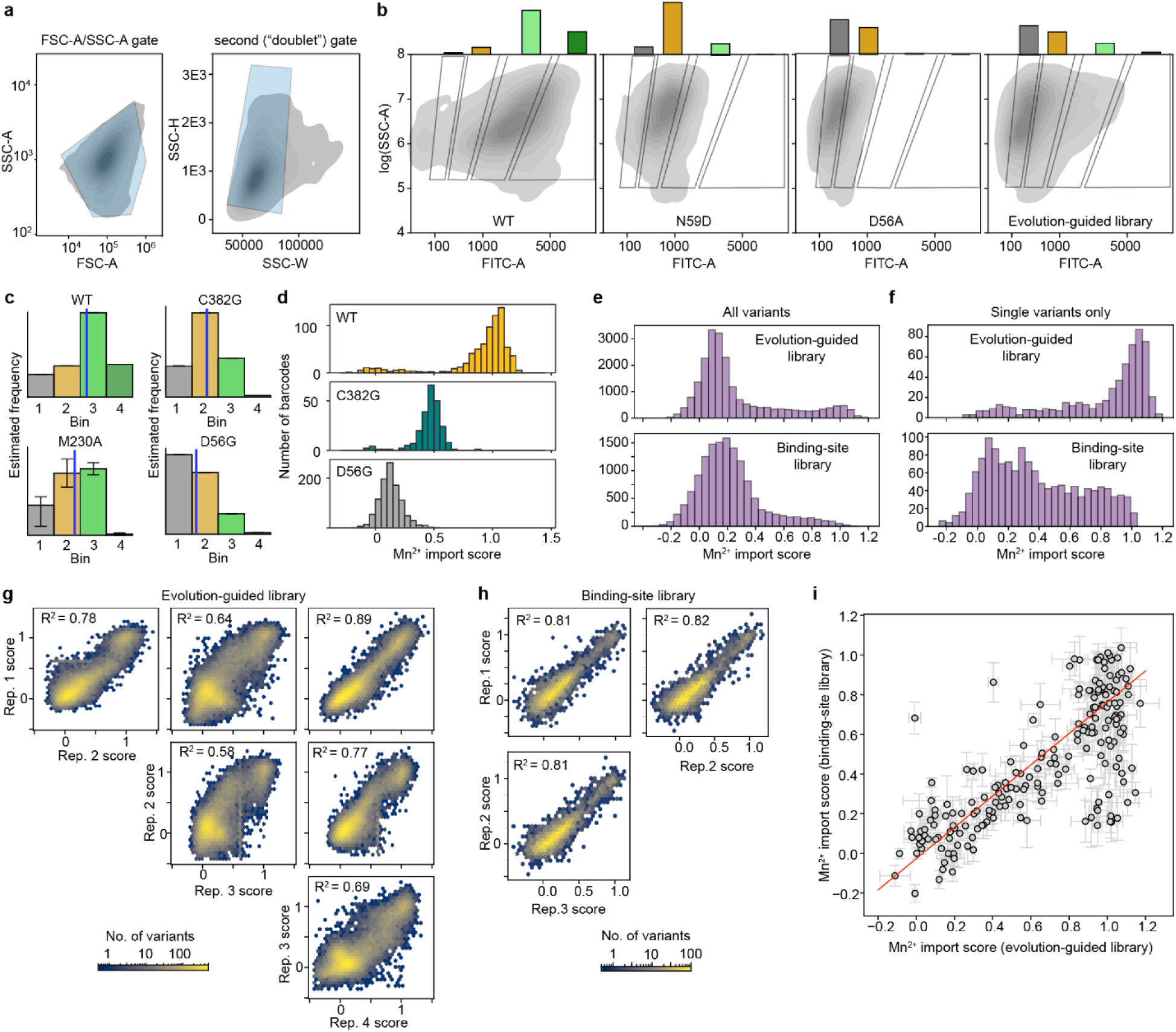
High-throughput screen for Mn^2+^ import. (a) Example flow cytometry gating scheme, from the first replicate of the evolution-guided library, with the overall cell population is shown as a grey density plot and the gates in transparent blue. The FSC-A/SSC-A gate (left) largely selects against rare events outside of the core distribution of cells. The second (“doublet”) gate (right) excludes events with unusual values of SSC-W compared to SSC-H, which are more likely to represent clusters of multiple cells. (b) Gating schemes for the four green fluorescence gates for sorting, shown with the cell population density plots for the three controls (DraNramp WT, DraNramp N59D, DraNramp D56A) and the first replicate of the evolution-guided library. Distributions of cells across the gates are shown as bars above their respective gates. (c) Example estimated frequency histograms for the 4 bins for 4 variants: WT, C382G, M230A, and D56G, all from the first replicate of the evolution-guided library. Error bars represent standard deviations from 100 resamples from *p*(*λ*|*c*_*i*_) (see *Methods*), which are visible only for M230A due to the smaller number of reads observed for M230A than the other example variants. Y-axes are unitless and thus labels are excluded. The blue vertical line represents the weighted mean bin value for that variant. (d) Distributions of Mn^2+^ import scores calculated across different barcodes representing the three most abundant variants in the evolution-guided library: WT (top), C382G (middle), and D56G (bottom). (e) Distribution of scores for all variants in the evolution-guided and binding-site libraries, pre-merging. (f) Distribution of all scores for single substitution variants in each library. (g-h) Replicate correlations between individual replicates in the evolution-guided (g) and binding-site (h) libraries. All replicates correlate well other than replicate 3 of the evolution-guided library, which still correlates with the others but somewhat worse, indicating more experimental error in the collection of this replicate. (i) Correlation of replicates for variants overlapping between the two libraries, with regression line overlaid in red. Error bars on points represent estimated errors based on both replicate- and read-count based error.

**Extended Data Figure 4.**
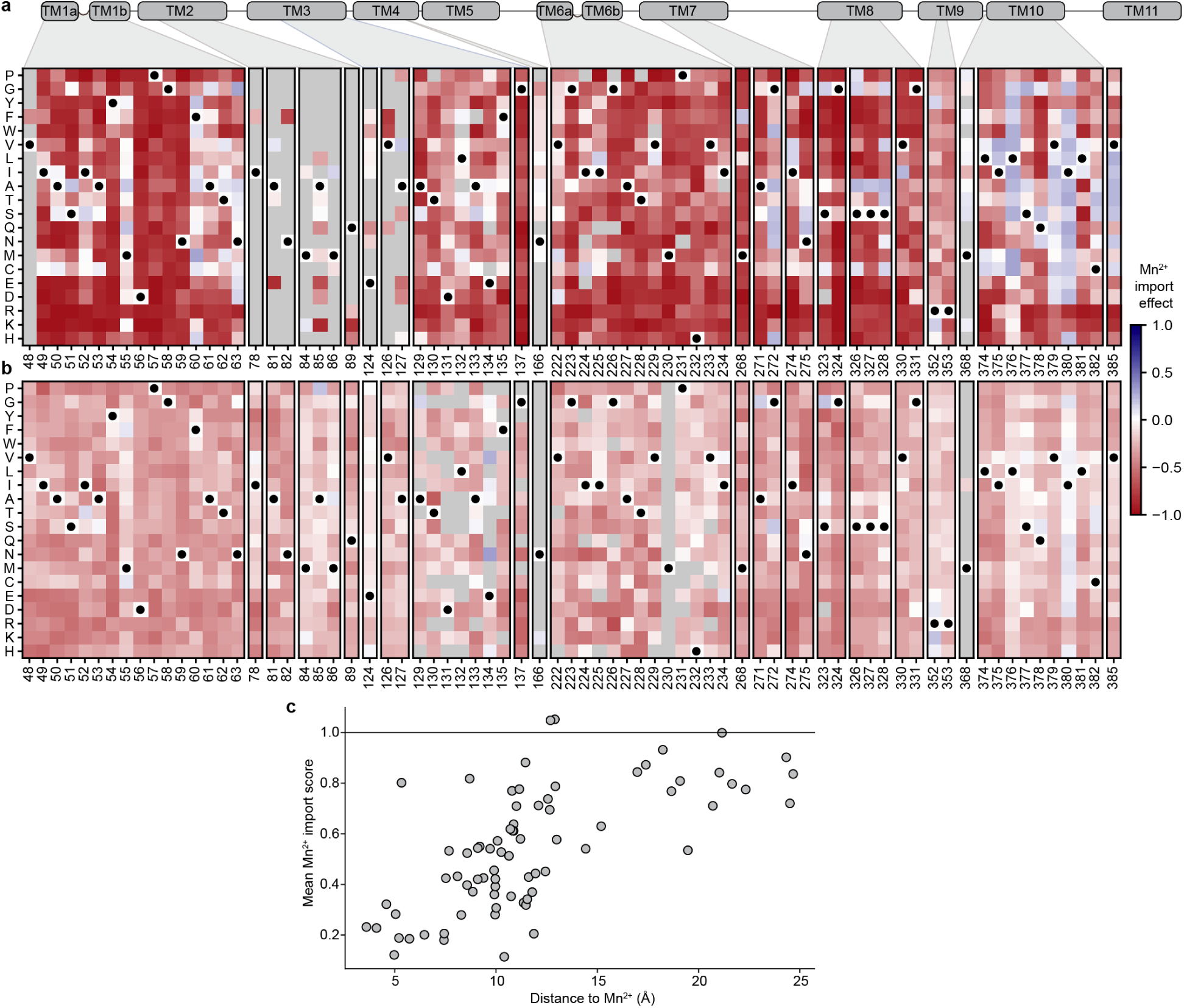
Mutation effects on Mn^2+^ import score. (a) Heatmap showing the effect of each single mutation in the combined library dataset, obtained by subtracting the WT score of 1 from the corresponding variant score, such that WT DraNramp has a score of 0. Positions with at least seven variants in either background are shown. The TM2 region is largely absent from the WT-background library due to an unknown error in assembly of the binding-site library from individual subpools, thus only positions represented in the evolution-guided library have data. (b) Analogous heatmap showing the effect of each single mutation on the M230A background, such that M230A has an effect of 0. (c) Plot of mean Mn^2+^ import score for variants with single mutations at a particular residue position as a function of that residue’s distance from the bound Mn^2+^ substrate.

**Extended Data Figure 5.**
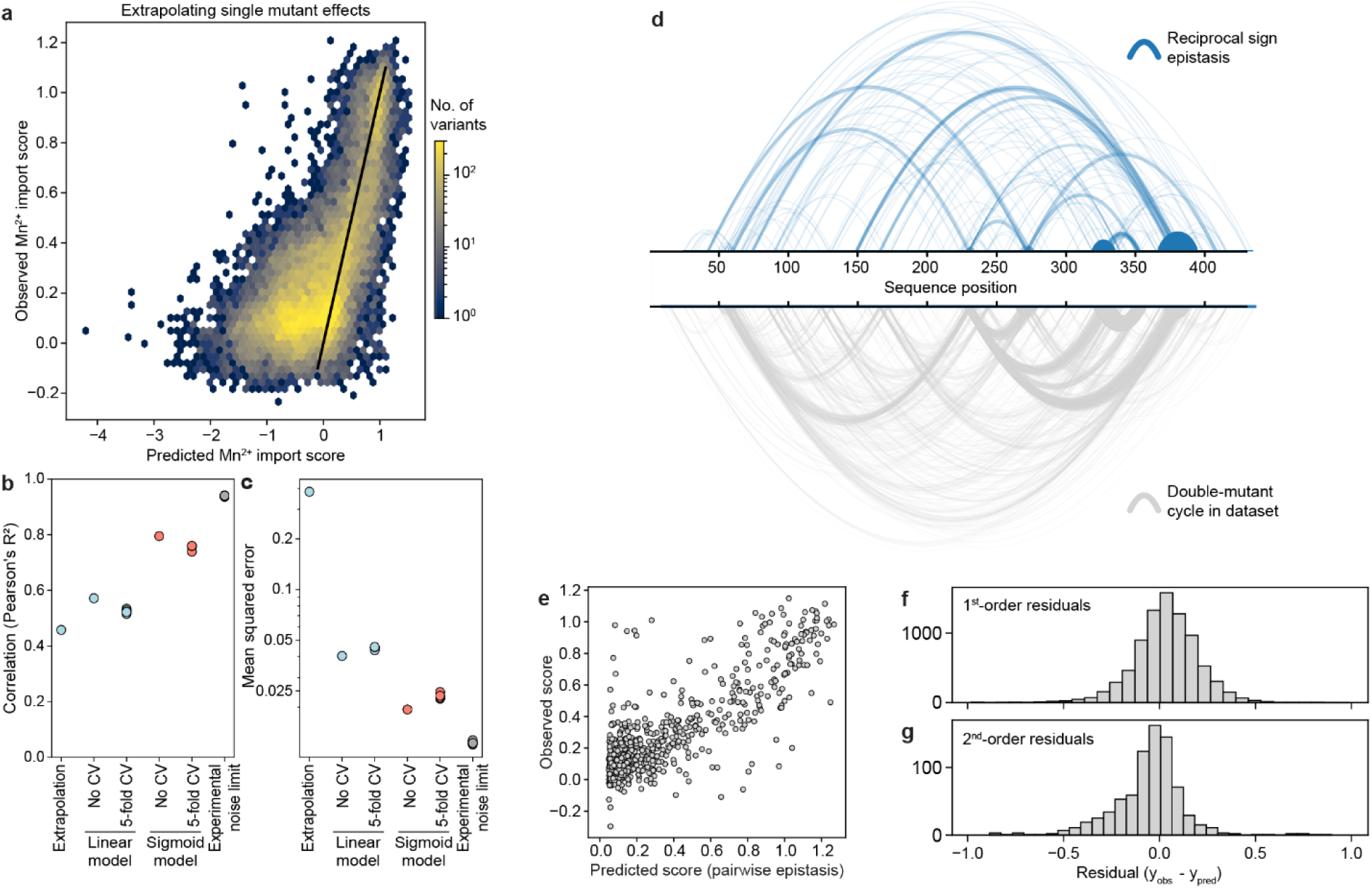
Additional analyses related to modeling the Mn^2+^ fitness landscape. (a) Density plot comparing predicted and observed Mn^2+^ import scores for all variants with two or more mutations by extrapolating the mutation effects from the single variants. A black linke indicates perfect prediction. (b) Summary of the Pearson’s correlation coefficients for the predicted Mn^2+^ import score of variants against their measured values with five models: a simple linear extrapolation of single mutant effects to higher-order variants (panel a), or linear regression (Fig. 3a) or sigmoidal regression (Fig. 3b) models incorporating global epistasis, each reported with or without five-fold cross-validation (CV), with the correlations for the five held-out folds shown independently for the relevant columns. The gray point represents the theoretical maximum correlation achievable from random samples from the experimental error. (c) An equivalent plot showing the mean squared error (MSE) of the prediction compared to the measured value. Note that the y-axis is on a logarithmic scale to capture the extremely high MSE of the extrapolation model without obscuring other differences. (d) Arc plot showing pairs of positions with reciprocal sign epistasis (blue) compared against all pairs present in the dataset (gray). In each case, the frequency of each double mutant in the two sets is represented by the width and opacity of the line. (e) Predicted mutation effects incorporating couplings terms for each double-mutant cycle into the sigmoid regression model, evaluated on triple-mutant cycles. (f) Distribution of 1^st^-order residuals among double-mutant cycles (similar to Figure 3c but for exclusively double mutants). (g) Distribution of 2^nd^-order residuals among triple-mutant cycles incorporating pairwise epistasis terms.

**Extended Data Figure 6.**
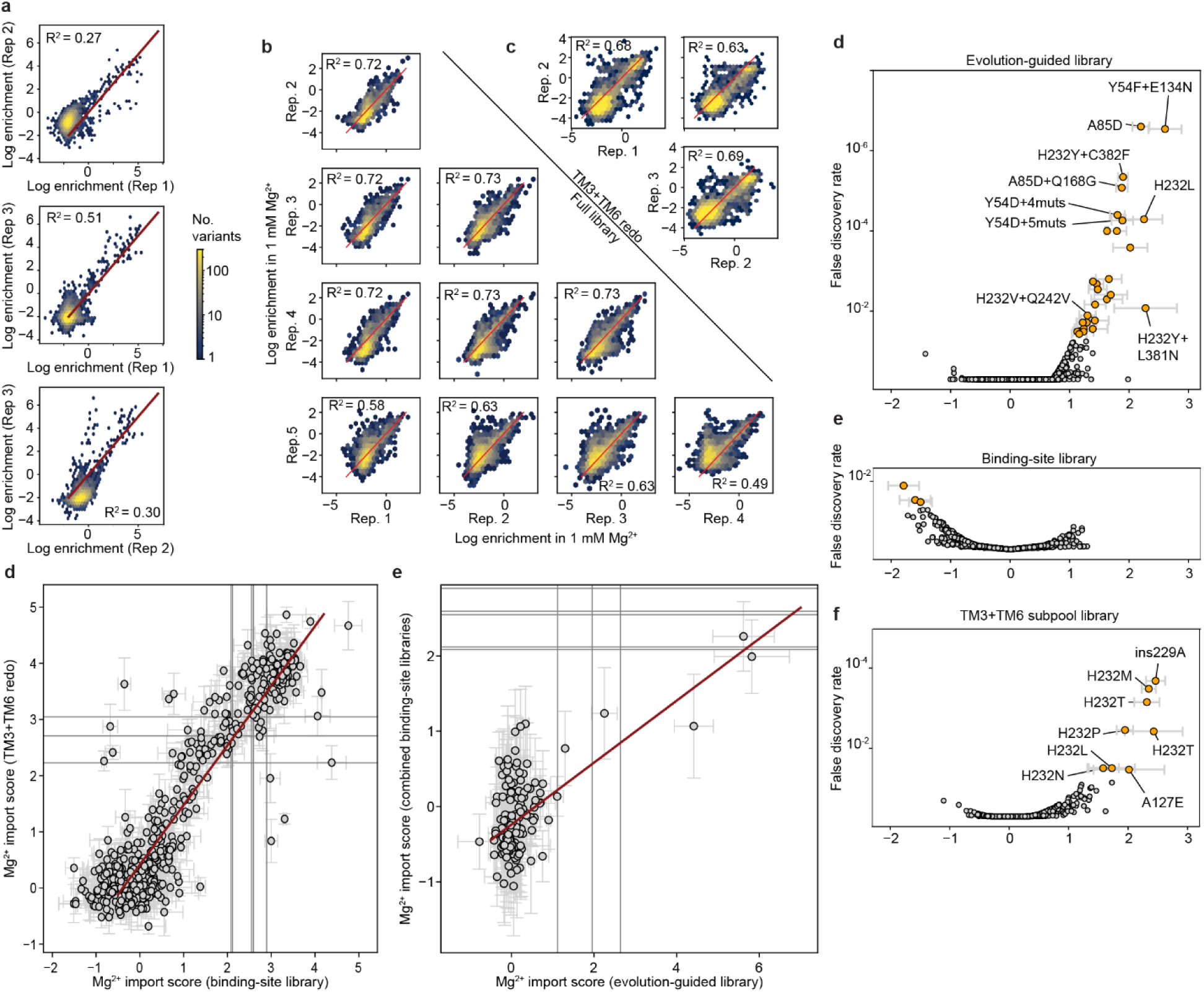
Statistical analysis of Mg^2+^ screen results. (a-c) Density plots showing correlation between each pair of replicates of the Mg^2+^ screen for the evolution-guided library (a), the full binding-site library (b), and the repeat of the screen with only the TM3 and TM6 subpools (c). Replicate scores generally correlate well, albeit with generally lower Pearson’s R^2^ values than in the Mn^2+^ import screen (ranging from 0.27 to 0.73 across 16 comparisons). Note that the highly skewed population distribution of the evolution-guided library in particular, where most variants have scores within noise of zero, explains the lower observed correlation compared to many deep mutational screen libraries testing proteins’ native activities. (d) Orthogonal distance regression plot showing the correlation of overlapping Mg^2+^ import scores between the full binding-site library screen (x-axis) and the TM3+TM6 subpool screen (y-axis), with Pearson’s R^2^ = 0.81. Error bars on points represent estimated experimental error, and red line represents fit regression line. Vertical and horizontal gray bars show the scores for mutants with no enrichment after growth in 1mM MgSO_4_ (raw log enrichment score of zero before normalization to WT), one bar per replicate. In both libraries, most variants were significantly depleted (left of or below gray bars); only a few were enriched (right of or above gray bars). (e) Similar regression plot for the combined binding-site library against the evolution-guided library, with Pearson’s R^2^ = 0.51. In this case, the pre-normalization zero point scores (gray) are very different. (f-h) Volcano plots comparing the raw Mg^2+^ import score and the false discovery rate for M230-retaining variants in the evolution-guided library (f), the full binding-site library (g), or the TM3+TM6 subpool library (h). False discovery rates were calculated based on the null distribution of barcodes mapping to WT DraNramp, the estimated error on each experimental observation, and then adjusted for multiple hypotheses with the Bejamini-Hochberg correction (see Methods). In orange are the 37 M230-retaining variants that are significantly enriched.

**Extended Data Figure 7.**
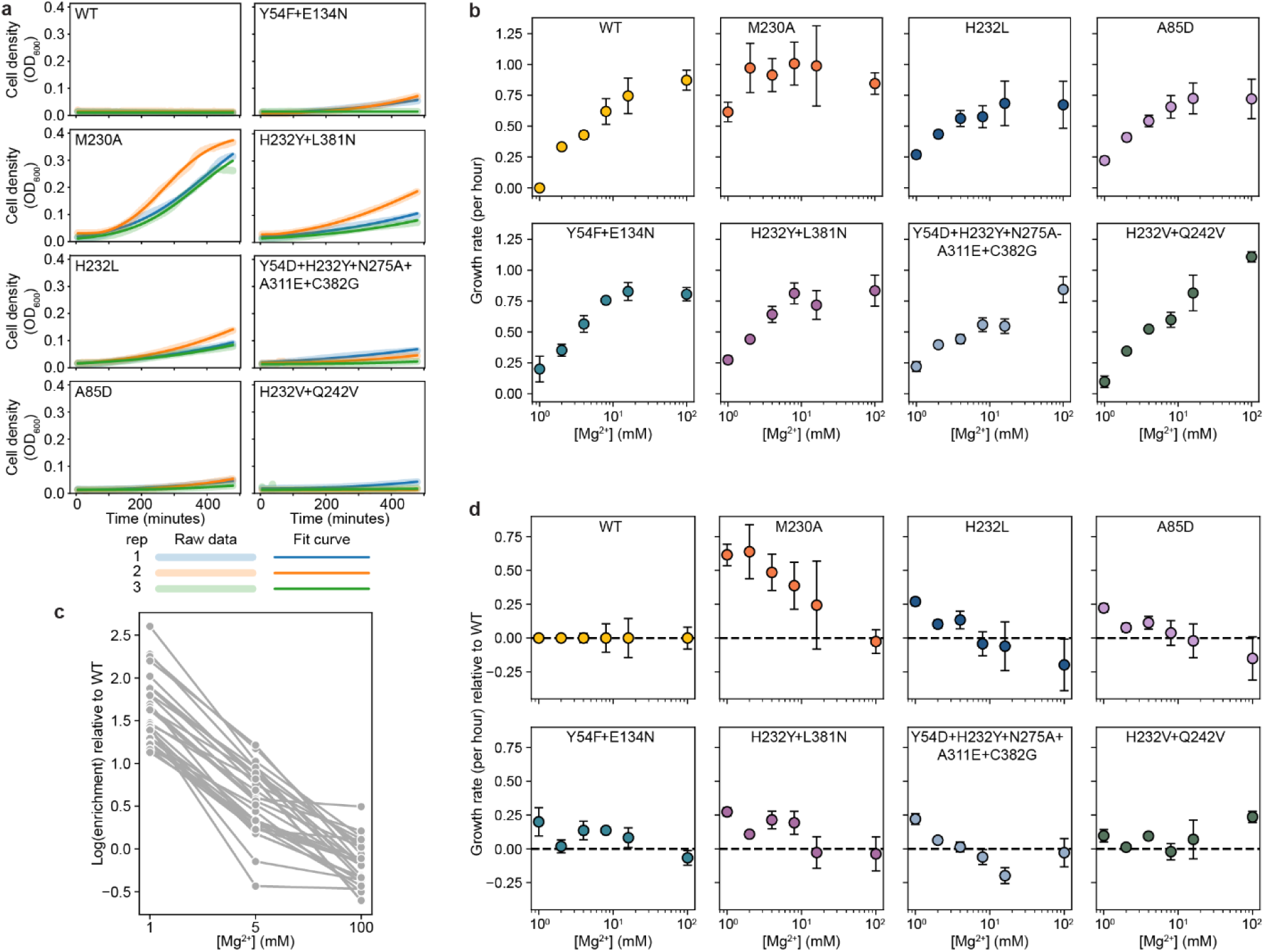
Growth curves and dose responses for DraNramp variants in the Mg^2+^ growth complementation assay. (a) Growth curves for DraNramp variants tested individually at 1mM MgSO_4_, with each of three independent biological replicates colored differently. The fit growth curve, from which the growth rate was then extracted, is overlaid as a thinner opaque line. (b) Growth rates of different DraNramp variants as a function of Mg^2+^ concentration. Error bars at each point represent the standard error of the mean. (c) Log-enrichment scores relative to WT for all 37 M230-containing variants with significant Mg^2+^ import scores in the selective 1 mM condition (see Extended Data Fig. 6f-h) across three concentrations of Mg^2+^: a stringent 1-mM selection, a weak 5-mM selection, and 100 mM, with no selection on Mg^2+^ import ability. All variants have a clear negative trend, showing that their ability to improve growth relative to WT scales with the strength of the selection. (d) The same growth rates for variants tested in isolation as in (b), normalized to the growth rate of WT at that [Mg^2+^] to allow for comparison to the wild-type protein as in (c). Variants that rescue growth by transporting Mg^2+^— rather than through other means—would be expected to have an improvement in growth over WT that scales with the stringency of the Mg^2+^ selection, which we observe for all of the tested variants and especially M230A, H232L, and Y54D+H232Y+N275A+A311E+C328G.

**Extended Data Figure 8.**
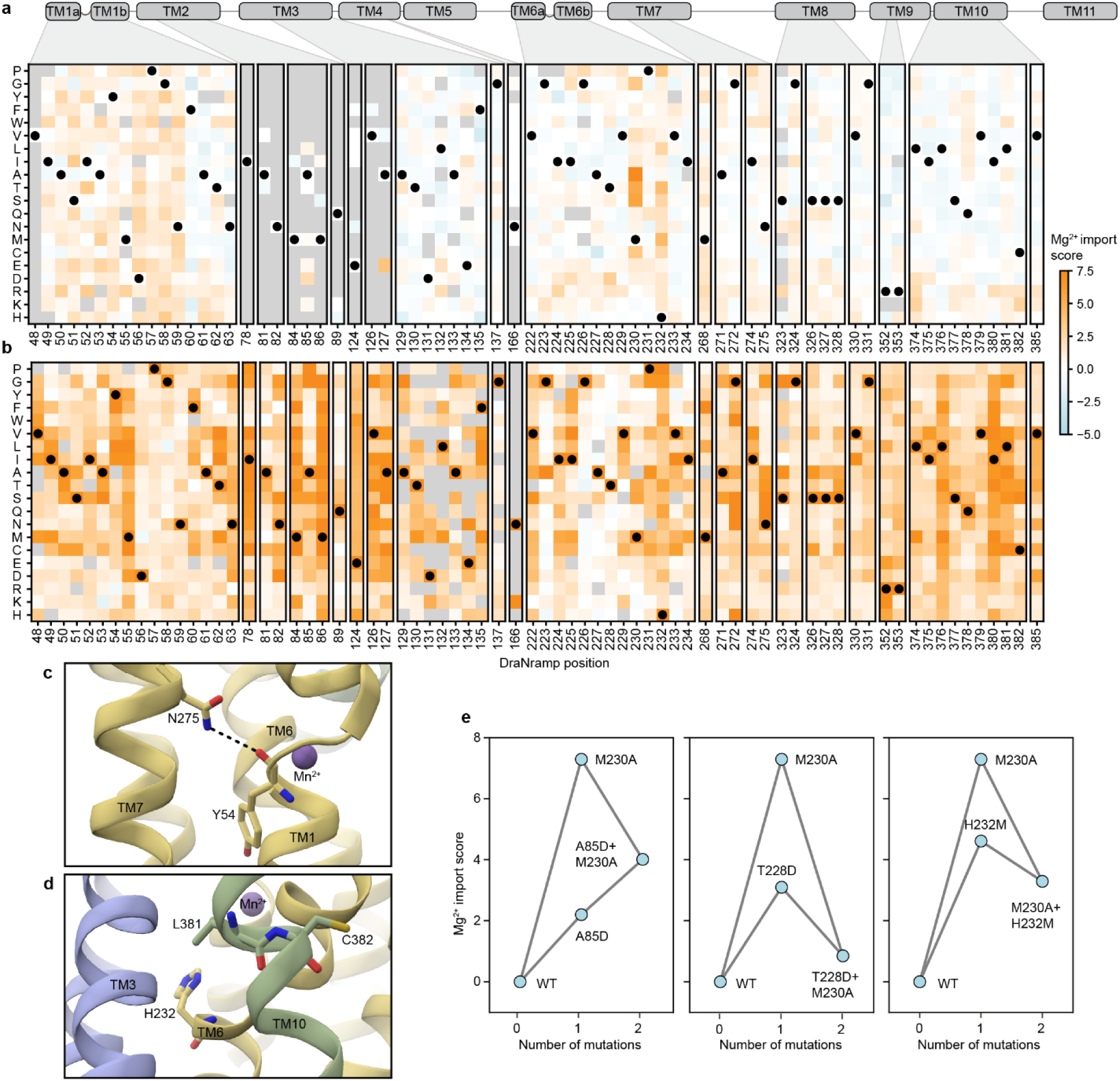
A map of Mg^2+^ import in the WT and M230A backgrounds. (a-b) Heatmaps of Mg^2+^ import scores for residues with at least 10 mutations on either the WT (a) or M230A (b) backgrounds. Black circles mark wildtype amino acids. Gray positions lack data. (c-d) Structural snapshot from the DraNramp occluded structure (PDB ID: 8E60) illustrating the proximity of Y54 and N275 (c) and H232, L381, and C382 (d). (e) Three double-mutant cycle plots illustrating examples of negative sign epistasis for the A85D, T228D and H232M mutations on the WT vs. M230A backgrounds.

**Extended Data Figure 9.**
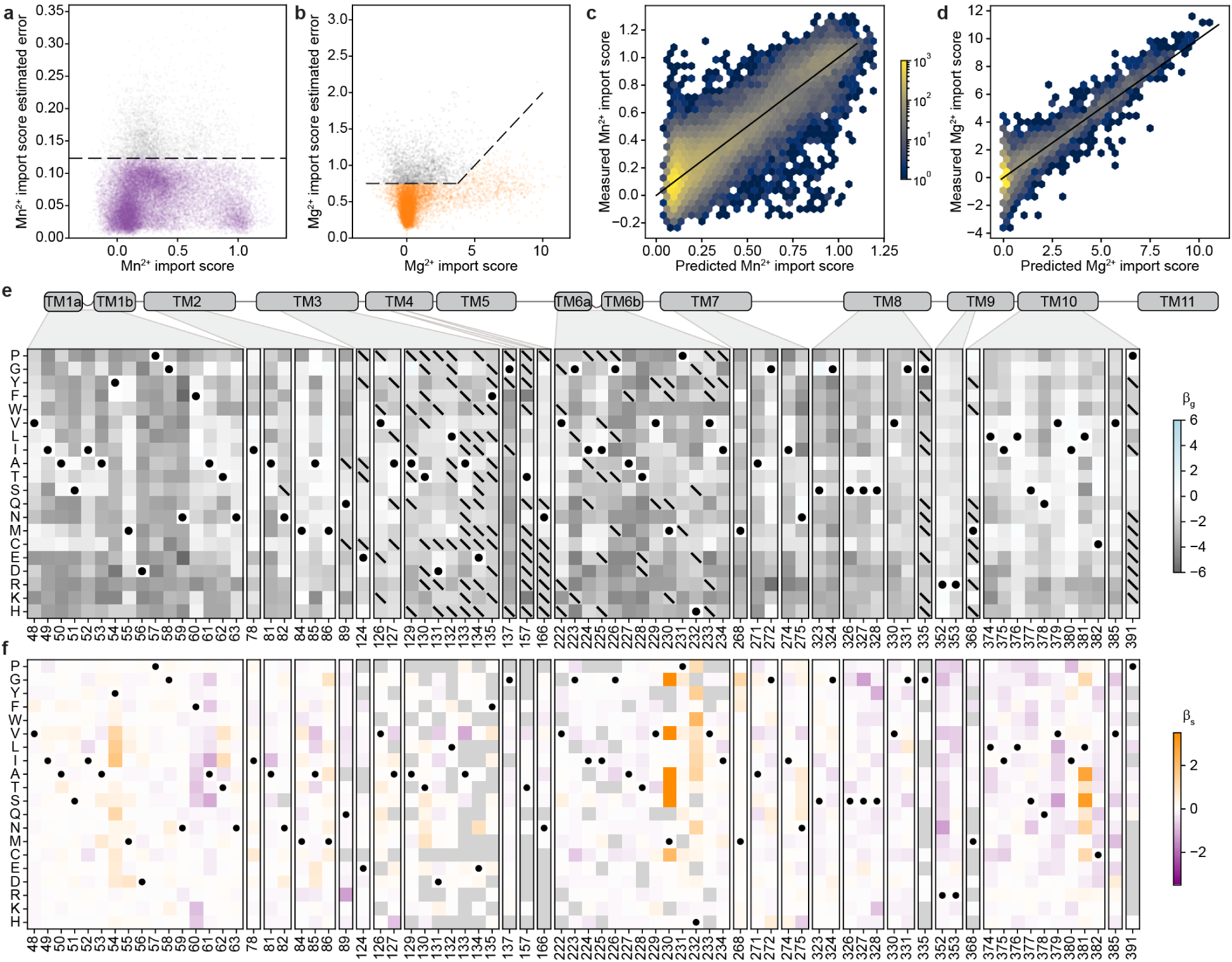
Additional analysis associated with Figure 5. (a) The Mn^2+^ import score and associated estimated error for each variant in the combined library, with the error cutoff shown as a dashed line. (b) The same for Mg^2+^, highlighting how the error cutoff increases for larger Mg^2+^ score values. (c) Predicted Mn^2+^ import score against measured score for all variants in the unmasked sigmoid specificity model, without cross-validation as we are using this model purely as a descriptive model, not a predictive one. (d) Predicted Mg^2+^ import score against measured score for all variants in the unmasked sigmoid specificity model, as in (c). (e) Posterior mean estimates for each *β*_*g*_ term in the combined specificity model, in logits. Terms that were not fit are marked with a diagonal line. (f) Posterior mean estimates for each *β*_*s*_ term in the combined specificity model, with positive terms (orange) representing mutations that shift the protein toward Mg^2+^ specificity and negative terms (purple) those that shift the protein toward Mn^2+^ specificity. Terms that were not fit are light gray.

