## Supplemental Table 1 and Supplementary Notes 1-2 for "Determinants of metal import and specificity in a bacterial transporter"

This file contains:

- Supplementary Table 1. Comparison of activity parameters inferred from $Mn^{2+}$ import assay in bacteria to those determined *in vitro*
- Supplementary Note 1. Estimation of minimal fitness landscape models with stochastic variational inference
- Supplementary Note 2. Derivation and interpretation of conformational balance curve
- Supplementary Figures 1-5

The following tables are provided as separate CSV or Excel documents:

- Supplementary Table 2. All processed $Mn^{2+}$ and $Mg^{2+}$ import scores.
- Supplementary Table 3. Variants passing significance thresholds for growth rescue in the $Mg^{2+}$ assay.
- Supplementary Table 4. Binding-site library oligonucleotide pools.
- Supplementary Table 5. Design for the spread-out low-diversity library used to build the evolution-guided library.
- Supplementary Table 6. All primers used in this study, including primers for cloning individual DraNramp variants, primers used for preparing Illumina sequencing, and primers for assembling the binding-site library.

**Supplementary Table 1.** Comparison of activity parameters inferred from $Mn^{2+}$ import assay in bacteria to those determined *in vitro*

| Variant | Kinetic parameters from data in Figure 2c | | Data from the literature[a] | | | |
|---|---|---|---|---|---|---|
| | $\widetilde{K_{m,eff}}$ (µM) | Relative $\widetilde{V_{max}}$ | Membrane potential (mV) | External pH | $K_m$ (µM) | $V_{max}$ (µM /min) |
| WT | 22 ± 3 | 1 ± 0.03 | -50 | 7 | 700 ± 100 | 20 ± 2 |
| | | | -50 | 6 | 54 ± 3 | 34.1 ± 0.5 |
| | | | -100 | 7 | 7.0 ± 0.8 | 21.6 ± 0.5 |
| | | | -100 | 6 | 4.4 ± 0.3 | 37.2 ± 0.6 |
| | | | -120 | 7 | 7 ± 3[b] | 20 ± 1[b] |
| | | | -150 | 7 | 1.8 ± 0.2<br>3.4 ± 0.2[c] | 28.1 ± 0.6<br>54 ± 1[c] |
| | | | -150 | 6 | 2.9 ± 0.2 | 40.8 ± 0.6 |
| D56A | N/A | 0 | 0 to -120 | 5.7 to 7 | N/A | 0 |
| M230A | 12.4 ± 3.9[d] | 0.61 ± 0.06[d] | -150 | 7 | 97 ± 8 | 50 ± 1 |
| | | | -150 | 7 | 140 ± 10[c] | 83 ± 2[c] |
| N59D | 5.6 ± 1.5 | 0.39 ± 0.03 | -150 | 7 | 38 ± 2[c] | 10.8 ± 0.1[c] |

[a]Except where noted, the values are from [1].

[b]Values from [2].

[c]Values from [3].

[d]*In vivo* dose-response curves for M230A do not saturate (Fig. 2C); thus, the $\widetilde{K_{m,eff}}$ and $\widetilde{V_{max}}$ from a simple fit are likely underestimated. We cannot reliably test saturating concentrations of $MnCl_2$ for M230A as concentrations greater than 400 µM begin to significantly inhibit cell growth (with many cells appearing dead in FACS at $MnCl_2$ concentrations of 400 µM or higher), so accurately estimating these parameters is challenging. This means that the higher $V_{max}$ and $K_m$ estimated from *in vitro* studies testing $Mn^{2+}$ concentrations up to 1 mM [1,3] shown above are consistent with our results. In comparing our values from *in vivo* estimates to those from *in vitro* experiments under controlled buffer conditions, it is worth noting that the effects of competing $Mg^{2+}$ and $Ca^{2+}$ in the medium are difficult to predict, as they could competitively inhibit $Mn^{2+}$ transport but could also noncompetitively compete by binding and stabilizing transport-inactive states such as the inward-open conformation.

**Supplementary Note 1. Estimation of minimal fitness landscape models with stochastic variational inference**

To better understand how mutations combine, we fit a set of regression models to the DraNramp fitness landscape. Following other models incorporating global epistasis[4-7], we make the fundamental assumption that the observed phenotype $y_i$ is a nonlinear function of an underlying latent factor (or fitness potential) $z_i$, and that these unobserved latent factors have a simpler additive relationship than the actual observed phenotypes. The major difference is that we estimate a posterior distribution over these latent factors using stochastic variational inference as implemented in pyro[8]. This allows us to explicitly incorporate structured priors into our inference framework and learn confidence distributions over our inferred parameters.

The following code is implemented on top of pyro v1.9.1[8] in the python package flambé v0.1 (fitness landscape architecture models with Bayesian estimation), which is currently in development.

*Linear model*

We begin with a straightforward linear regression model which does not include a latent variable $z_i$ (Supplementary Fig. 1). This model assumes the following likelihood function:

$$p(y_i|\theta_i) \sim Normal(x_i^T \beta + b, \sigma_i^2)$$

Here, $x_i$ is a one-hot encoded vector of the individual mutations present the variant $i$, with wildtype amino acid identities removed. $\beta$ is a vector of the regression weights (or variant effects) for each mutation in the dataset. The intercept $b$ is added to all variants and thus represents the wildtype activity level. Finally, the variance of this distribution is estimated as a combination of the experimental variance estimated on each variant, $\sigma_{obs}^2$, with an additional model variance term $\sigma_{model}^2$ that accounts for additional irreducible variance stemming from the inaccurate linear approximation of the landscape:

$$\sigma_i^2 = \sigma_{obs}^2 + \sigma_m^2$$

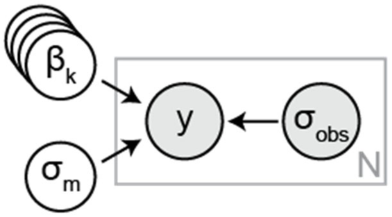


**Supplementary Figure 1.** Stylized graphical representation of a linear regression model. Random variables in white circles are inferred by the model, while gray circles ($y$ and $\sigma_{obs}$) represent directly observed data. The light gray box represents a "plate" in pyro's nomeclature, demonstrating that there is a "copy" of the data within it for each data point in the training set. By contrast, we use stacked circles for the parameters $\beta_i$ to represent how this parameter has a term for each feature but lies outside of the plate (so is constant for each data point).

We set the following priors on the parameters:

$$\beta_k \sim Normal(0.,2.)$$

$$b \sim Normal(0.,2.)$$

$$\sigma_m \sim HalfNormal(0.1)$$

The relatively broad Gaussian priors centered around zero represent a generally uninformative prior that makes no assumptions on the average direction of mutational effects but does assume that they will not be extraordinarily large in magnitude. The halfnormal prior on $\sigma_m$ is similarly relatively uninformative but constrains the error to be positive and not huge.

We then constructed a mean-field variational posterior (or "guide") $q_\phi$ with a normal distribution on on $\beta$ and $b$ and a lognormal distribution on $\sigma_m$, as the error cannot be negative. The assumption of normality for each posterior distribution will lead to somewhat inaccurate error estimates, but it is much simpler than alternatives and more computationally efficient. Relaxation of these assumptions to more accurately model dependencies will be a project for future study.

To train this model, pyro's automated stochastic variational inference (SVI) routine then optimizes the parameters of the variational posterior $q_\phi$ to maximize the evidence-based lower bound (ELBO), given by the difference between the expected value of the likelihood of the data given the model and the deviation from the prior:

$$L(\phi) = E_{q_\phi}[\log(p(y|X,\theta)] - KL(q_\phi||p(\theta))$$

To efficiently train this model, we use a ClippedAdam optimizer, starting with a high learning rate (0.2) that decays by multiplying by a factor of 0.99 per iteration over 3,000 iterations.

*Site-independent sigmoid model*

The next simplest model is the site-independent sigmoid model (Supplementary Fig. 2). This is the first model that assumes the existence of a latent variable $z_i$:

$$z_i = x_i^T \beta + b$$

$$\hat{y}_i = k + \frac{a}{1 + \exp(z_i)}$$

$$p(y_i|\theta_i) \sim Normal(\hat{y}, \sigma_i^2)$$

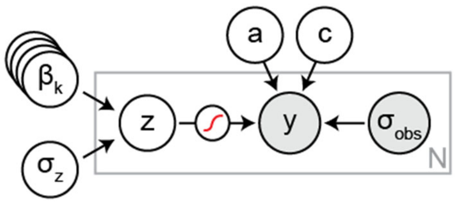


**Supplementary Figure 2.** Graphical representation of the site-independent sigmoid model.

Here, $a$ and $k$ are inferred scalar parameters that form part of the sigmoid transformation to rescale the predicted score from (0,1) to the natural range of the data. We also assign Gaussian priors to these parameters. For the $Mn^{2+}$ dataset, they are:

$$a_{\mathrm{Mn}} \sim Normal(1.,0.5)$$

$$k \sim Normal(0.,0.5)$$

These priors are slightly more informative than those on $\beta$ and $k$ because for these data we have a strong prior intuition as to the shape of the sigmoid. In particular, without any prior or constraint on $a$, all of the model's parameters could flip in sign; here, however, we have a strong prior that $a$ is more likely to be positive. We also know to expect a particular scale. This means that the prior on $a$ should be adjusted for datasets with very different scales. For the $Mg^{2+}$ dataset, which has generally larger values due to the nature of the data processing, we change the prior on $a$ to:

$$a_{\mathrm{Mg}} \sim Normal(2.,5.)$$

We represent this model with the stylized graphical model in Supplementary Figure 3. After training this model, we want to be able to invert the sigmoidal transformation to estimate $p(z_i|y_i)$. One challenge is that some observations $y_{obs,i}$ are outside of the range of the sigmoidal function. We will assume that this is due to noise rather than true biological variability and thus $k < \hat{y} < k + a$. By Bayes' rule,

$$p(\widehat{y_i}|y_{obs,i}) = p(y_{obs,i}|\widehat{y_i})p(\widehat{y_i})$$

Although many forms could be assumed for the prior distribution of a particular output $p(\widehat{y_i})$, for simplicity we assume that it is uniform within the range $(k, k + a)$. As $p(y_{obs,i}|\widehat{y_i}) \sim Norm(\widehat{y_i})$, we can therefore estimate $p(\widehat{y_i}|y_{obs,i})$ as a truncated normal distribution $\psi(y_{obs,i}, \sigma_{obs,i}, k, a + k)$. Therefore, to sample from $p(z_i|y_i)$, we first sample a set of parameters from the posterior guide, then sample $\widehat{y_i}$ from $\psi(y_{obs,i}, \sigma_{obs,i}, k, a + k)$, and calculate:

$$z_i = \log(\frac{a}{\widehat{y_i} - k}) - 1$$

*Two-substrate independent sigmoid model*

We further adapt this model to integrate both datasets together (Supplementary Fig. 3). In this case, we aim to share information between datasets by building in the assumption that most effects will be shared between data. To this end, we first define a global latent activity:

$$z_{g,i} = x_i^T \beta_g$$

We can then add a new set of specific terms to this:

$$z_i = z_{g,i} + x_i^T \beta_s$$

While in theory we could have a separate set of terms for each substrate, with only two substrates this is not very meaningful; to avoid overfitting uninterpretable parameters we do not fit specificity terms for the $Mn^{2+}$ import assay data and thus define $z_s$ as the difference between manganese and magnesium import, as follows:

$$\hat{y}_{\mathrm{Mn}} = k_{\mathrm{Mn}} + \frac{a_{\mathrm{Mn}}}{1 + \exp(z_{g,i} + b_{\mathrm{Mn}})}$$

$$\hat{y}_{\mathrm{Mg}} = k_{\mathrm{Mg}} + \frac{a_{\mathrm{Mg}}}{1 + \exp(z_{g,i} + z_{s,i} + b_{\mathrm{Mg}})}$$

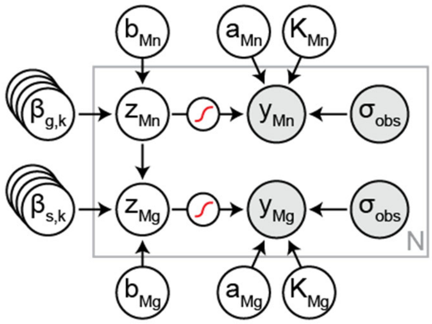


**Supplementary Figure 3.** Stylized graphical representation of the two-substrate sigmoid model.

Our initial assumption motivating the joint model is that mutations directly affecting specificity are rarer than those affecting global parameters of the protein (and thus affecting manganese and magnesium import similarly). Without any such assumption, the joint model is quite similar to simply fitting two separate models. To demonstrate this, imagine fitting a separate sigmoid for each substrate with parameters $\beta_{Mn}$ and $\beta_{Mg}$. The likelihood term of the ELBO for the joint model would simply be the sum of the likelihoods for these independent models, and thus the model would be driven to fit the same parameters, except that $\beta_s$ would be $\beta_{Mg} - \beta_{Mn}$. However, the prior terms can be different. We implement the assumption that relatively few terms affect specificity directly by placing a sparse Laplace prior on $\beta_s$, which will drive unnecessary $\beta_s$ to zero and push the model to explain as much of the variance with $\beta_g$ as possible (see Figure 6a).

One additional detail of this model is the ability to mask certain data points or parameters. We wanted to be able to learn from data points present in either dataset without requiring them to be in both. Therefore, the likelihood function for the specificity model can accept a mask representing missing data, and the likelihood is only evaluated over unmasked points. Similarly, we allow for fixing certain terms in $\beta_s$ to zero by masking them out during inference as well. For example, we fit a model that assumes that only position 230 contributes to specificity by masking out all features in $\beta_s$ that do not correspond to mutations at position 230.

**Supplementary note 2. Derivation and interpretation of the conformational balance curve**

The following note contains the derivation for and further explanation of Equation 1 (the conformational balance curve).

*Derivation*

We sought to develop a simple model of the effect on $V_{max}$ of mutations that alter the energetic balance between the inward- and outward-open states of the importer under facilitated diffusion conditions. Importantly, we will assume that the mutation affects only this and no other biochemical parameters of the protein. The overall reaction can be modeled as a cycle as in Supplementary Figure 4.

Here, $V_{max}$ represents the rate of increase of substrate on the inside ($\frac{d[S_{in}]}{dt}$) when $S_{in} = 0$ and $S_{out} \to \infty$. Under these conditions, the reaction is purely driven forward, and all equilibria are effectively irreversible, giving the simpler reaction scheme shown in Supplementary Figure 4.

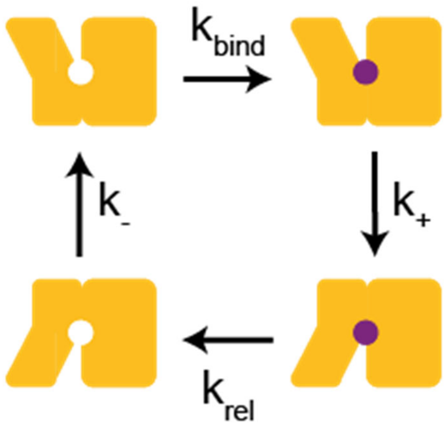


**Supplementary Figure 4.** Simple reaction cycle for import by facilitated diffusion.

To obtain $\frac{d[S_{in}]}{dt}$, we are interested in the total flux through this cyclic reaction pathway, which under a steady-state assumption[9] is given by the harmonic mean of the individual rates:

$$V_{max} = \frac{[T]}{\frac{1}{k_{bind}} + \frac{1}{k_+} + \frac{1}{k_{release}} + \frac{1}{k_-}}$$

where $[T]$ is the total concentration of the correctly folded transporter in all conformations.

To simplify this assumption, we make the classic enzyme kinetics assumption that substrate binding and release are much slower than the "catalytic" step, which in this case is transporter flipping, so we are left with:

$$V_{max} = [T]\left(\frac{1}{k_+} + \frac{1}{k_-}\right)^{-1}$$

Using the Arrhenius equation,

$$k_+ = B_{out}\exp(\frac{-\Delta G^{\ddagger}_{out\to in}}{RT})$$

and

$$k_- = B_{in}\exp(\frac{-\Delta G^{\ddagger}_{in\to out}}{RT})$$

Importantly, $\Delta G^{\ddagger}_{in\to out}$ and $\Delta G^{\ddagger}_{out\to in}$ are the transition states of similar reactions in opposite directions. If a mutation perturbs $\Delta G_{out\to in}$, it must influence either one or likely both of $\Delta G^{\ddagger}_{in\to out}$ and $\Delta G^{\ddagger}_{out\to in}$. Following $\phi$ value theory, we parameterize this relationship with $0 < \phi < 1$, such that:

$$\Delta\Delta G^{\ddagger}_{out\to in} = \phi\Delta\Delta G_{out\to in}$$

$$\Delta\Delta G^{\ddagger}_{in\to out} = (1-\phi)\Delta\Delta G_{in\to out}$$

Because $\Delta\Delta G_{in\to out} = -\Delta\Delta G_{out\to in}$ we can also write:

$$\Delta G^{\ddagger}_{in\to out} = (\phi - 1)\Delta\Delta G_{out\to in}$$

For simplicity, we will from now on refer to the variable $\frac{\Delta\Delta G_{out\to in}}{RT}$, as $x$.

Therefore, upon mutation, we have a set of new forward and reverse rates:

$$k'_+ = B_{out}\exp(-\frac{\Delta G^{\ddagger}_{out\to in} + \Delta\Delta G^{\ddagger}_{out\to in}}{RT})$$

$$= B_{out}\exp(-\frac{\Delta G^{\ddagger}_{out\to in}}{RT} + x)$$

$$k'_- = B_{in}\exp(-\frac{\Delta G^{\ddagger}_{in\to out}}{RT} + (\phi - 1)x)$$

These can be simplified by expressing the new rates after mutation in terms of the original rates $k^0_+$ and $k^0_-$:

$$k'_+ = k^0_+ e^{-\phi x}$$

$$k'_- = k^0_- e^{-(\phi-1)x}$$

Substituting these terms back into the original flux equation, we have:

$$V'_{max} = [T]\left(\frac{e^{\phi x}}{k^0_+} + \frac{e^{(\phi-1)x}}{k^0_-}\right)^{-1}$$

At this point, we observe that this defines a nonlinear equation of $\Delta\Delta G_{out\to in}$, with $\phi$, $[T]$, $k^0_+$ and $k^0_-$ as constants. To better understand its behavior, we separate terms that rescale the whole function from those that change its shape. To do so, it is useful to extract a factor of $\frac{1}{k^0_+}$ from the equation:

$$V'_{max} = \frac{[T]}{k_+^0}\left(e^{\phi x} + \frac{k_+^0}{k_-^0} e^{(\phi-1)x)}\right)^{-1}$$

Here, the term $\frac{[T]}{k_+^0}$ does not affect the shape of the curve, only its magnitude, while the shape depends only on $\phi$ and the ratio of the initial forward and reverse rates (or the "pseudo-equilibrium constant") $\frac{k_+^0}{k_-^0}$. In the case where the substrate does not affect the rate of the forward reaction at all, this will be the baseline equilibrium constant between the two states. However, in practice, this will not be the case for an efficient transporter as the substrate likely stimulates the forward reaction and $\frac{k_+^0}{k_-^0}$ will be larger than $K_{eq}$.

Finally, to better understand the effect of a mutation, we will write out this equation in terms of the original $V_{max}^0$, which is given by:

$$V_{max} = \frac{[T]}{k_+^0}\left(1 + \frac{k_+^0}{k_-^0}\right)^{-1}$$

Thus:

$$\frac{V'_{max}}{V_{max}^0} = \left(1 + \frac{k_+^0}{k_-^0}\right)\left(e^{\phi x} + \frac{k_+^0}{k_-^0} e^{(\phi-1)x}\right)^{-1}$$

This is equivalent to Equation 1 in the main text.

*Interpretation*

The equation above describes a family of curves parameterized by $\phi$ and $\frac{k_+^0}{k_-^0}$. The very simplest family member would have $\phi = \frac{1}{2}$, indicating a fully symmetric transition state, and $\frac{k_+^0}{k_-^0} = 1$ , indicating that the forward and reverse reactions happen at the same rate. In this case, the equation will simplify to:

$$\frac{V'_{max}}{V_{max}^0} = 2\left(e^{\frac{x}{2}} + e^{-\frac{x}{2}}\right)^{-1} = \text{sech}(\frac{x}{2})$$

The simplest family member is thus equivalent to the well-studied hyperbolic secant curve. Understanding this also helps us see how changing $\frac{k_+^0}{k_-^0}$ changes the curve's behavior. For simplicity, below we define $C = \frac{k_+^0}{k_-^0}$. Then we rewrite:

$$e^{\frac{x}{2}} + Ce^{-\frac{x}{2}} = \sqrt{C}\left(e^{\frac{x-\ln C}{2}} + e^{-(\frac{x-\ln C}{2})}\right) = \frac{\sqrt{C}}{2}\cosh(x - \frac{\ln C}{2})$$

Therefore:

$$\frac{V'_{max}}{V^0_{max}} = \frac{1+C}{\sqrt{C}} \text{sech}(\frac{x}{2} - \frac{\ln C}{2})$$

Note that this simply represents a linear transformation of the original hyperbolic secant curve, with the position of the maximum shifted by $\frac{\ln C}{2}$ and the overall height of the curve scaled by $\frac{1+C}{\sqrt{C}}$. This both provides an analytical solution to how the optimal $\Delta\Delta G_{out\rightarrow in}$ changes with $\frac{k^0_+}{k^0_-}$ and shows that varying $\frac{k^0_+}{k^0_-}$ does not alter the overall shape of the curve when $\phi = \frac{1}{2}$.

For an asymmetric transition state with $\phi \neq \frac{1}{2}$, the curve can no longer be described by a hyperbolic secant (Supplementary Fig. 5). Thus, $\phi$ does fundamentally change the shape of the curve, unlike $\frac{k^0_+}{k^0_-}$.

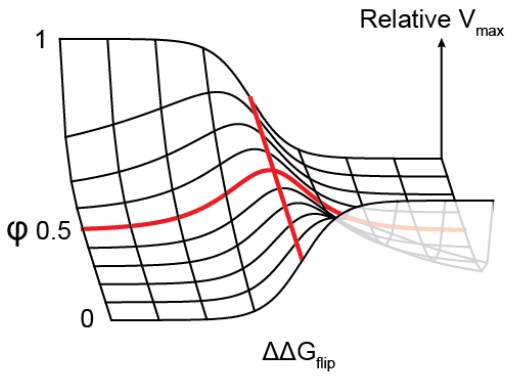


**Supplementary Figure 5.** Changes in the conformational balance curve as the constant parametrizing the linear free energy relationship, $\phi$, changes. This fundamentally changes the shape of the curve, unlike the $\frac{k^0_+}{k^0_-}$ parameter (see Figure 7c).
